# CYRI-B mediated macropinocytosis drives metastasis via lysophosphatidic acid receptor uptake

**DOI:** 10.1101/2022.11.23.517689

**Authors:** Savvas Nikolaou, Amelie Juin, Jamie A. Whitelaw, Nikki R. Paul, Loic Fort, Colin Nixon, Heather J. Spence, Sheila Bryson, Laura M. Machesky

**Author notes:** **Conflict of interest:** The authors declare that they have no potential conflicts of interest.

## Abstract

Pancreatic ductal adenocarcinoma carries a dismal prognosis, with high rates of metastasis and few treatment options. Hyperactivation of KRAS in almost all tumours drives RAC1 activation, conferring enhanced migratory and proliferative capacity as well as macropinocytosis. Macropinocytosis is well understood as a nutrient scavenging mechanism, but little is known about its functions in trafficking of signaling receptors. We find that CYRI-B is highly expressed in pancreatic tumours in a mouse model of KRAS and p53- driven pancreatic cancer. Deletion of CYRI-B accelerates tumourigenesis, leading to enhanced ERK and JNK-induced proliferation in precancerous lesions, indicating a role as a buffer of RAC1 hyperactivation in early stages. However, as disease progresses, loss of CYRI-B inhibits metastasis. CYRI-B depleted tumour cells show reduced chemotactic responses to lysophosphatidic acid, a major driver of tumour spread, due to impaired macropinocytic uptake of LPAR1 receptor. Overall, we implicate CYRI-B as a mediator of growth and signaling in pancreatic cancer, providing new insights into pathways controlling metastasis.

## Introduction

Pancreatic ductal adenocarcinoma (PDAC) is highly metastatic with low survival rates and few treatment options. PDAC is thought to arise from precancerous non-invasive pancreatic intraepithelial neoplasms (PanINs) classified as PanIN1-3 depending on the molecular and histological characteristics (Hruban *et al*, 2001; Hruban *et al*, 2007). PanINs arise from acinar cells that undergo acinar to ductal metaplasia changes (Wang *et al*, 2019) and as mutations accrue, PanINs progress to full PDAC in which angiogenesis, infiltration of stromal cells and invasion of the basement membrane occur as tumours progress. Metastasis is a complex process and the current gold standard mouse model of metastatic PDAC -the KPC (KRAS^G12D^, p53^R172H^, Pdx-1::Cre) (Hingorani *et al*, 2005) recapitulates multiple features of the human disease (Hwang *et al*, 2016). Cytoskeletal and migration-associated proteins have been associated with aggression and metastasis in PDAC both in human patient transcriptomes (Bailey *et al*, 2016) and in the KPC mouse model (Juin *et al*, 2019; Li *et al*, 2014) suggesting avenues to pursue against metastatic spread.

Downstream of active KRAS, hyperactivation of the small GTPase RAC1 drives proliferation and cytoskeletal remodeling in PDAC and other cancers. Deletion of RAC1 in a KRAS-driven mouse model of PDAC delayed tumour onset, reduced PanIN lesions and improved survival (Heid *et al*, 2011; Wu *et al*, 2014). This led to the conclusion that dysregulation of RAC1 control of epithelial polarity by active KRAS drives acinar to ductal metaplasia and accelerates tumourigenesis (Heid *et al*., 2011). RAC1 regulates polarity and migration via Scar/WAVE- Arp2/3 control of actin dynamics at cell-cell contacts and at the cell leading edge. Additionally, coordinated RAC1 activation and deactivation are important in macropinocytosis, an actin-driven process whereby cells engulf extracellular substances via large cup-shaped protrusions of the plasma membrane (Egami *et al*, 2014). Extracellular stimulation of cell surface receptors, such as tyrosine kinase or G-protein coupled receptors, can trigger macropinocytosis via RAC1 and the Scar/WAVE complex (Buckley & King, 2017). Tumours are frequently starved for amino acids and other nutrients and macropinocytosis is a major way for PDAC tumours to take up proteins, lipids and cell debris from their environment (Commisso *et al*, 2013; Hobbs & Der, 2022; Kamphorst *et al*, 2015; Puccini *et al*, 2022; Yao *et al*, 2019). Macropinocytosis also provides cells with a mechanism for internalisation of signalling receptors (Clayton & Cousin, 2009; Le *et al*, 2021; Stow *et al*, 2020), but whether this has consequences for tumour progression is unknown.

Metastasis is a complex process, involving cells breaching through tissue barriers, migrating and settling in distant sites in the body such as the liver, lungs and peritoneal cavity (Nikolaou & Machesky, 2020). Chemotaxis is thought to be a key driver of metastasis and pancreatic cancer cells migrate towards lysophosphatidic acid (LPA) both in vitro and in vivo, contributing to metastasis (Juin *et al*., 2019; Papalazarou *et al*, 2020). LPA is a serum-derived chemotactic factor and was previously found to be consumed by melanoma and PDAC cells creating self-generated gradients contributing to metastasis (Juin *et al*., 2019; Muinonen-Martin *et al*, 2014).

Recently, the CYRI-B protein (**<u>Cy</u>**fip-related **<u>R</u>**AC1-**i**nteracting protein B, formerly known as Fam49-B) was discovered to interact with RAC1 and enhance leading edge actin dynamics by negatively regulating activation of the Scar/WAVE complex (Fort *et al*, 2018). Scar/WAVE is a pentameric complex that interacts both with RAC1 and Arp2/3 complex and triggers actin assembly in lamellipodia (Insall & Machesky, 2009). Deletion of CYRI-B in cultured cells enhanced lamellipodia stability, but did not impair migration speed (Fort *et al*., 2018). CYRI proteins also play an important role in macropinocytosis, via the RAC1-Scar/WAVE pathway (Le *et al*., 2021). These roles, along with the previous implication of RAC1 signaling in early and later stages of PDAC (Heid *et al*., 2011; Wu *et al*., 2014) suggested potential involvement of CYRI in invasion and metastasis. Here we demonstrate that CYRI-B is highly expressed in PDAC and can contribute to PDAC development, progression and metastasis. We discover a role for CYRI-B in signaling that drives proliferation in early lesions. Later, during metastasis, we find that CYRI-B is required for chemotaxis toward LPA, implicating macropinocytic uptake of LPAR1 in PDAC metastasis. Our study highlights CYRI-B as a potentially interesting new target in PDAC progression and metastasis and further elucidates the molecular mechanisms underpinning metastatic spread.

## Results

### CYRI-B expression increases in precancerous lesions and PDAC

The *Cyri-b* gene resides on human chromosome 8q24, near c-Myc, and is frequently amplified in many types of cancer, including pancreatic cancer (Nikolaou & Machesky, 2020). High expression of CYRI-B correlates with poor outcome in many cancers (Li *et al*, 2021; Xu *et al*, 2022a), including in human pancreatic cancer. To further investigate a potential role for *cyri-b* in pancreatic cancer we first assessed the expression of CYRI-B in the KPC mouse model of metastatic PDAC (Hingorani *et al*., 2005). In this model, PanIN develops by around 10 weeks and progresses to later stages toward 15 weeks, with full blown PDAC appearing at this stage and mice reaching endpoint with a half time of median 150-200 days. Tissue samples from 6, 10, 15-week old KPC mice were processed for RNA in situ hybridisation (ISH). At 6 weeks, before appearance of PanIN, CYRI-B was not detected in the pancreas (Fig. 1). Cyri-b expression was detectable by 10 weeks, especially around PanIN lesions, which remained stable until 15-weeks of age (Fig.1). Endpoint tumours showed a significant increase in the levels of CYRI-B (Fig.1) suggesting that the KPC model was a good model for exploring the role of CYRI-B expression during PDAC progression.

**Figure 1.**
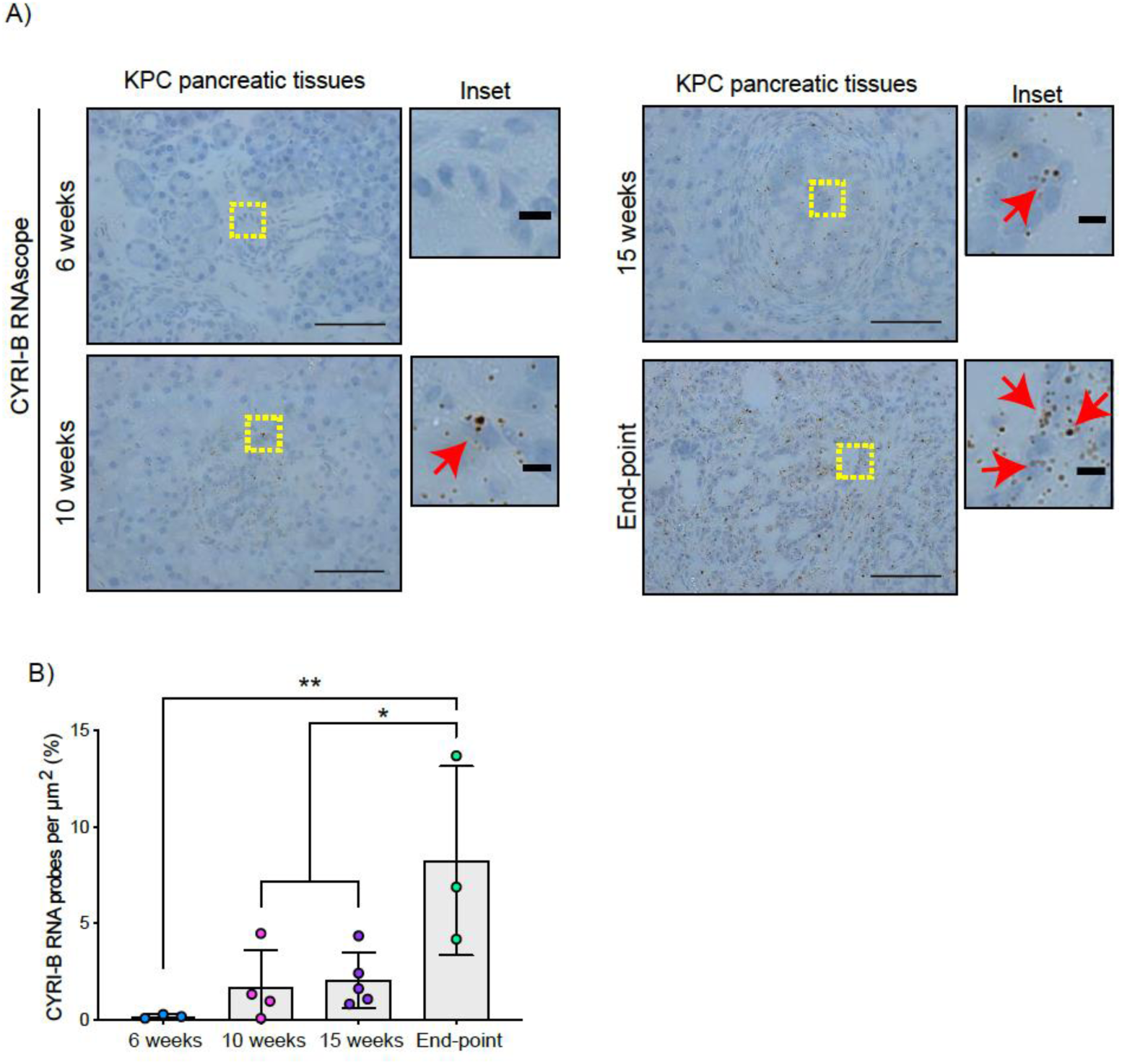
CYRI-B is expressed during PDAC progression. A) Representative images of CYRI-B RNA-scope in situ hybridisation from 6, 10, 15-week old and end-point KPC mouse tissues. RNA probes are visualised as brown dots. Haematoxylin was also used to stain the nuclei. Scale bars 50 mm. Yellow boxes show the region of interest for magnified images (inset). Red arrows denote positive RNA probes. Scale bars 5µm. B) Quantification of the CYRI-B RNA probes per mm^2^ from A). Mean ± SD; One-way ANOVA with Tukey’s test was performed in n≥3 mice. *p<0.01, **p<0.001.

### CYRI-B deletion accelerates PDAC development, reducing the survival of mice

To further probe the mechanism by which CYRI-B might influence PDAC progression, we crossed CYRI-B floxed mice with KPC mice. We refer to these mice and cell lines derived from them as CKPC (Fig. 2A). I*n situ* hybridisation (ISH) of end-point tumours confirmed no detectable *Cyri-b* mRNA in CKPC tumours in comparison with KPC (Fig. 2B,C). Western blotting also confirmed absence of CYRI-B protein in cell lines derived from end-point CKPC mouse tumors (CKPC-1 and CKPC-2) compared with a cell line from a KPC mouse tumor (KPC-1) (Fig. 2D). KPC and CKPC endpoint tumors showed no difference in the proliferation (Ki-67) or death (cleaved caspase-3, CC-3) of tumor cells (Fig. S1A-D). CKPC tumors also did not show any significant change in the CD31 vessel density (Fig. S1E-F) or necrosis (Fig. S1G-H), However, there was a significant decrease in the median survival to endpoint of CKPC mice (118 days) in comparison with the KPC mice (187 days) (Fig. 2E) without affecting the tumor weight to body mass ratio at end-point (Fig. 2F). Thus, loss of *Cyri-b* in the pancreas accelerates progression to endpoint of KRAS^G12D^, p53^R172H^ -driven PDAC in the KPC model, but does not grossly alter levels of cell growth/death or histological appearance of endpoint tumours.

**Figure 2.**
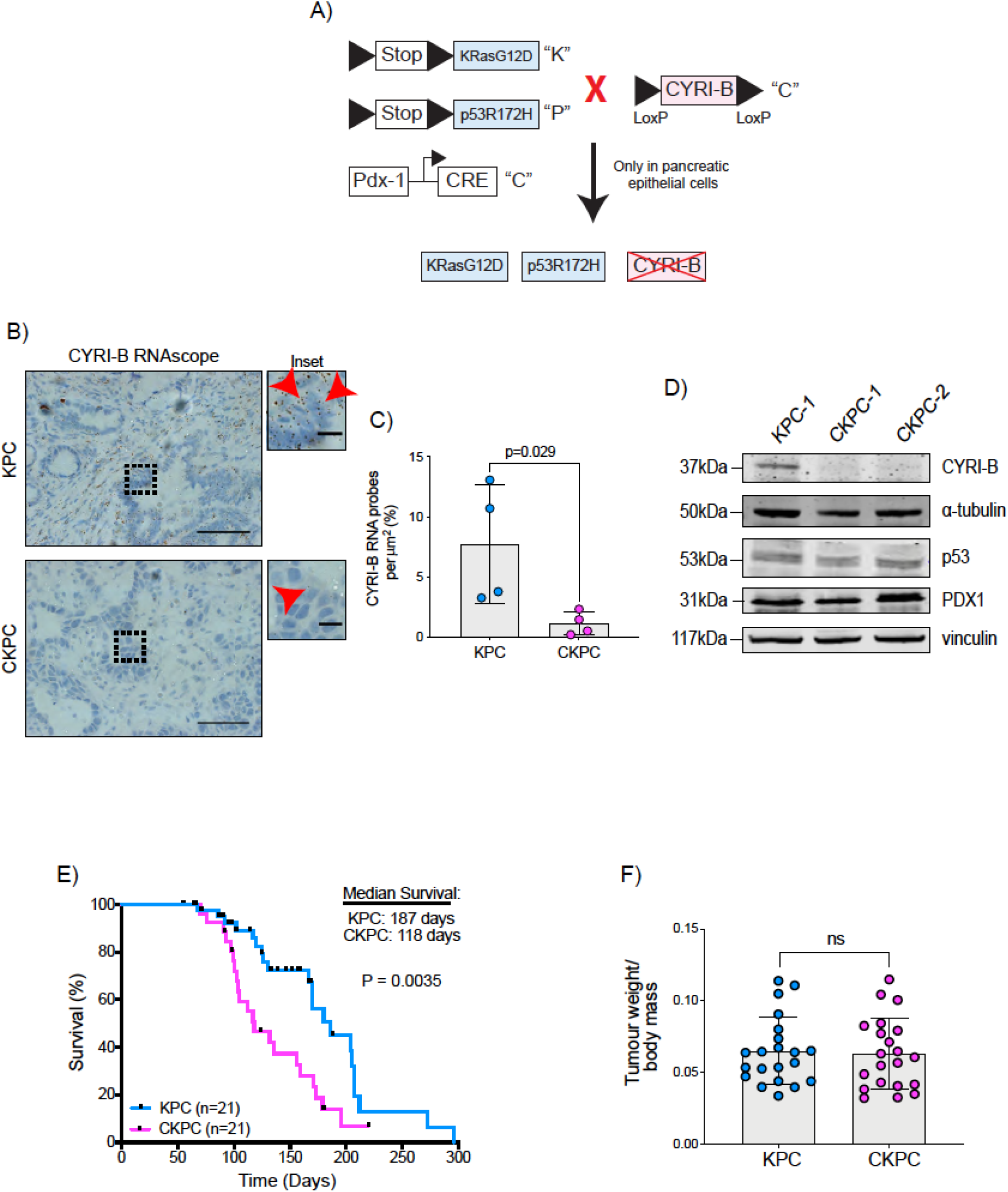
Loss of CYRI-B accelerates progression in the KPC mouse model of PDAC. A) Schematic representation of the CKPC mouse model. B) Representative images for CYRI-B RNA-scope staining of end-point tumours from KPC and CKPC mice. Scale bars, 50µm. Inset panels are magnified from the black dashed box. Scale bars, 10µm. Red arrows indicate the positive CYRI-B RNA. C) Histograms showing the CYRI-B RNA probes per mm^2^ at end-point tumours in KPC and CKPC mice. Mean ± SD; Unpaired t-test, n=4 KPC and 4 CKPC mice D) Representative Western blot images of CYRI-B in cell lines established from one KPC (KPC-1) and two CKPC (CKPC-1 and CKPC-2) tumours. Membranes were also probed for anti-p53 and anti-PDX1 to validate the CKPC cells. α-tubulin and vinculin were used as loading controls. Molecular weights as indicated on the side. See also Figure Supplement Source Data 1-5- original blots E) Survival (to endpoint) curve (n = 21 KPC, 21 CKPC independent mice). Log-rank (Mantel Cox) test used for comparing the KPC with CKPC survival curves. P-value as indicated. F) Histogram showing tumour-to-body mass ratios at sacrifice. Mean ± SD; Unpaired t-test was performed in n=21 KPC and 21 CKPC mice. p-value: not significant (ns).

### CYRI-B deletion accelerates PanIN formation

RAC1 is an important cancer driver downstream of KRAS and its ablation in mouse models delayed the onset of precancerous lesions (Heid *et al*., 2011) and led to an inability to sustain precancer progression to PDAC (Wu *et al*., 2014). Therefore, we asked whether loss of the RAC1 interactor CYRI-B affected the onset and progression between stages of PanIN1-3 (Fig. 3A,B). Pancreatic samples from KPC or CKPC mice revealed no differences at 10-weeks, but more PanIN2 and 3 lesions were present in 15-week old CKPC mice over KPC controls (Fig. 3C and Fig. S2), indicating an acceleration of early progression.

**Figure 3.**
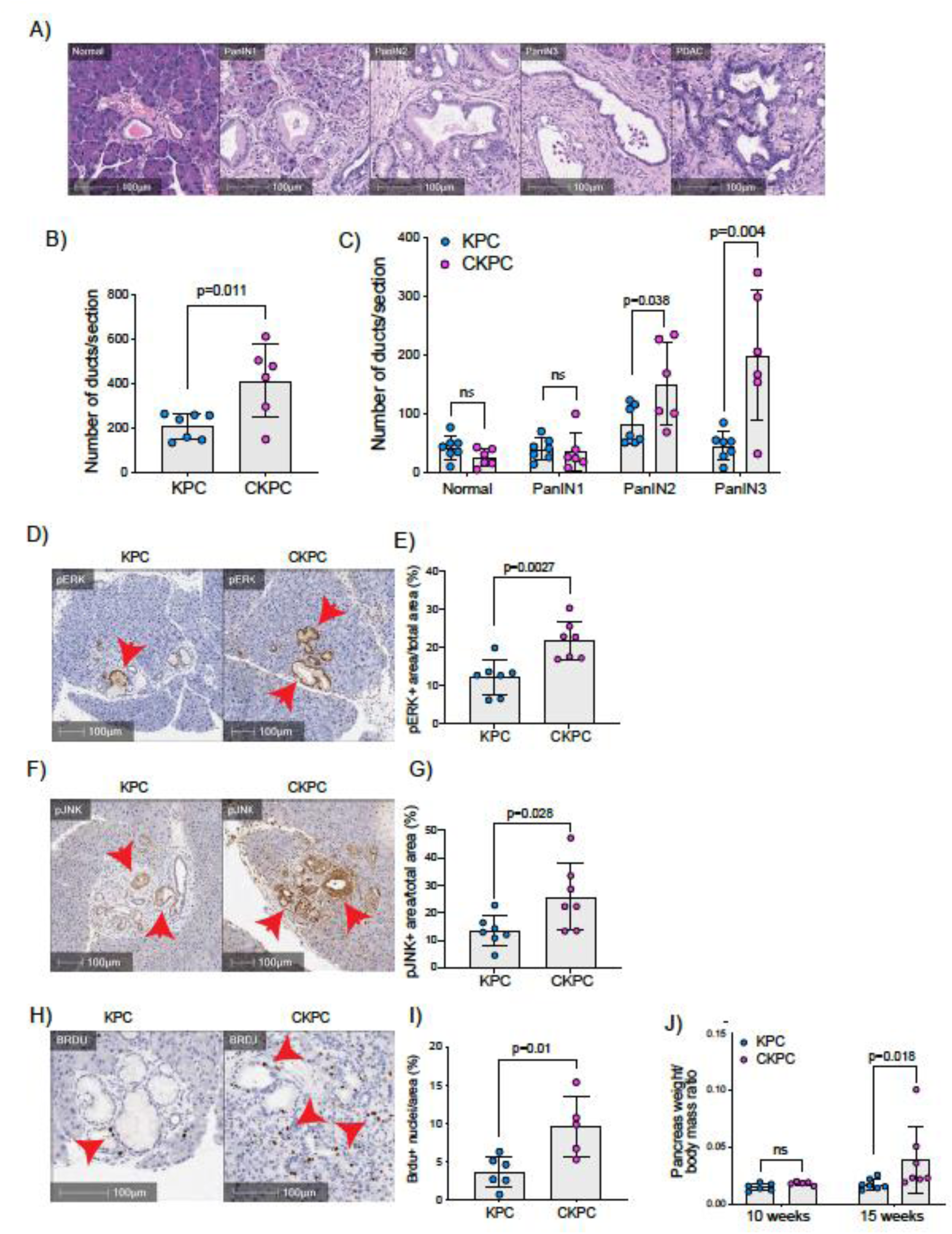
Loss of CYRI-B accelerates PanIN formation and increases pJNK, pERK and proliferation. A) Representative H&E images from KPC mice of normal pancreatic ducts, PanIN1, 2, 3 and PDAC lesions. Scale bars, 100µm. B) Number of ducts present in pancreas from 15-week-old KPC and CKPC mice (n≥6 mice). Mean ± SD; unpaired t-test was performed. p-value as indicated. C) Classification and scoring of pancreatic ducts in pancreas from 15-week-old KPC and CKPC mice (n≥6 mice). Mean ± SD; unpaired t-test was performed. ns=not significant, p-value as indicated. D) Representative images of pancreata from 15-week-old mice stained with pERK and haematoxylin (nuclei). Red arrows indicate the positive pERK staining. Scale bars, 100µm. E) pERK positive area from the total quantified area from D). Mean ± SD; Unpaired t-test was performed in n=7 KPC and CKPC independent mice. p-value as indicated. F) Representative images of pancreata from 15-week old mice stained with pJNK and haematoxylin (nuclei). Red arrows indicate the positive pJNK staining. Scale bars, 100µm. G) pJNK positive area from the total quantified area from F). Mean ± SD; Unpaired t-test was performed in n=7 KPC and CKPC independent mice. p-value as indicated. H) Representative images of 15-week-old KPC and CKPC mice stained for BrdU (proliferation) and haematoxylin. Red arrows show the BrdU positive nuclei. Scale bars, 100µm. I) Quantification of BrdU positive nuclei from KPC and CKPC 15-week-old pancreatic tissues. Mean ± SD; Unpaired t-test was performed in n=6 KPC and 5 CKPC independent mice. p-value as indicated. J) Quantification of the pancreas-to-body mass ratio at 10-weeks (n=6 mice in each mouse model) and 15-weeks (n=7 in each mouse model) in KPC and CKPC mice. Mean ± SD; Unpaired t-test was performed. ns=not significant, p-value as indicated.

To further probe the role of CYRI-B in lesion formation, we sought to understand the potential downstream signalling pathways that might be involved. RAC1 can drive cell proliferation through activation of both JNK and ERK downstream signalling pathways (Bagrodia *et al*, 1995; Coso *et al*, 1995; Rul *et al*, 2002; Wang *et al*, 2010). Therefore, we probed histological sections of pancreatic tissues from 15-week old KPC and CKPC mice for pJNK and pERK. Consistent with enhanced RAC1-signaling, we observed a significant increase in the percentage pERK area and pJNK area from pancreata of CKPC mice vs KPC (Fig. 3D-G). We next investigated proliferation using BrdU injections at 15 weeks and found increased BrdU positive nuclei in the CKPC tissues in comparison with the KPC suggesting enhanced proliferation in the abnormal ductal structures (Fig. 3H,I). Indeed, CKPC mice presented with increased pancreatic weight to body mass ratio at 15 weeks, in agreement with increased proliferation of preneoplastic and neoplastic cells, whereas at 10 weeks of age there was no difference (Fig. 3J). Thus, loss of CYRI-B in the KPC model accelerates PanIN formation and progression, likely due to loss of CYRI-B’s capacity to buffer RAC1 activation downstream of active KRAS leading to abnormal architecture, combined with hyperactivation of ERK and JNK to drive proliferation.

### CYRI-B regulates metastatic potential

The KPC mouse model is characterised by high metastatic rates to clinically relevant organs such as liver, diaphragm and bowel (Hingorani *et al*., 2005). Since CYRI-B regulates cell migration and chemotaxis (Fort *et al*., 2018), we asked whether deletion of CYRI-B can affect the metastatic potential of cancer cells in the CKPC mouse model. Analysis of mice at endpoint from KPC and CKPC cohorts revealed similar infrequent metastasis to the diaphragm in both cohorts, but a significant reduction in metastasis to both the liver and bowel of CKPC mice (Fig. 4A,B).

**Figure 4.**
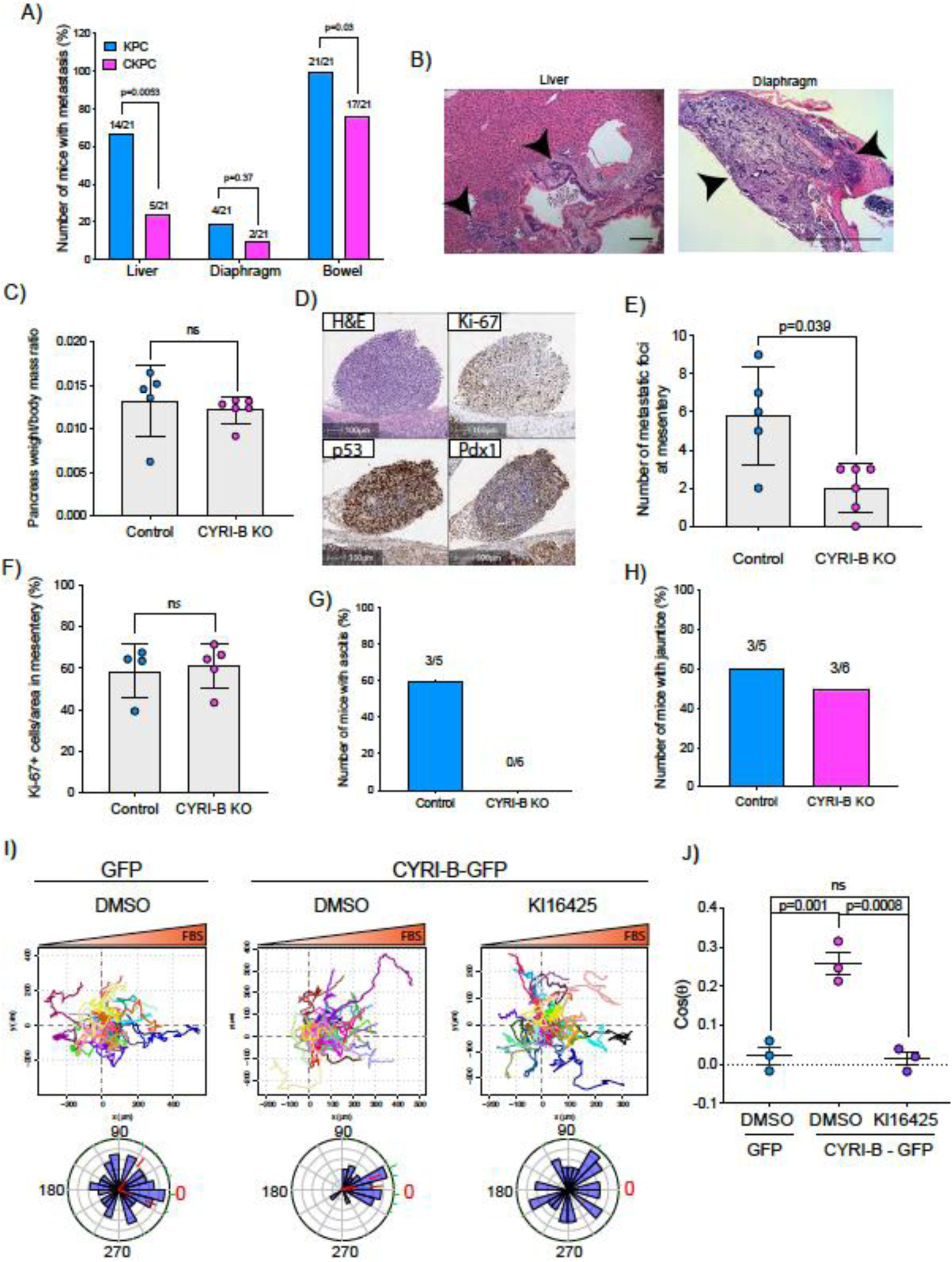
Loss of CYRI-B reduces metastasis and chemotactic potential. A) Incidence of KPC or CKPC mice presenting with metastasis in liver, diaphragm and bowel. Numbers above the bars show the fraction of mice with metastasis to the indicated site. Chi-square test was performed in n=21 KPC and CKPC mice. p-value as indicated. B) Representative H&E images of metastasis in the liver (Scale bar: 50µm) and diaphragm (Scale bar: 100µm). Black arrowheads denote metastatic lesions. C) Histogram showing pancreas-to-body mass ratios at sacrifice. Mean ± SD; Mann Whitney test was performed in n=5 for KPC control and n=6 mice for KPC CYRI-B KO cells. p-value: not significant (ns). D) Representative images of the mesenteric tumour foci from the in vivo transplantation assay. The metastatic foci were stained for haematoxylin and eosin, Ki-67 (proliferation), p53 and PDX1 (for control). Scale bars, 100µm. E) Histogram of the number of metastatic foci at mesentery for KPC control and KPC CYRI-B KO mice. Mean ± SD; Mann Whitney test t was performed in n≥5 mice for either control or CYRI-B KO KPC injected cells. p-value as indicated. F) Quantification of the Ki-67 positive cells in the metastatic tumour foci. Mean ± SD; Mann Whitney test was performed in n=4 for KPC control and n=5 mice for CYRI-B KO KPC cells. p-value: not significant (ns). G) Incidence of mice presenting ascites (n≥5). H) Incidence of mice presenting jaundice (n≥5). I) Representative spider-plots from n=3 independent chemotaxis assays of CKPC CYRI-B KO and rescued cells. A chemotactic gradient of 10% FBS was established and cells were imaged for 16h (1 frame/15min). Cells were also treated with either DMSO or the LPAR1/3 inhibitor KI16425 (10mM) for 1h prior to imaging. Each cell trajectory is displayed with a different colour and the displacement of each cell is reported in the x and y-axis. Orange gradient above shows the FBS gradient. Rose-plot data are displayed for each condition below. Red dashed lines show the 95% confidence interval for the mean direction in the Rose plots. The numbers represent degrees of the angle of migration relative to the chemoattractant gradient, with zero (red) denoting the direction of the chemoattractant gradient. J) Quantification of the results in I) showing the cos(θ) data (chemotactic index). Mean ± S.E.M from the average cos(θ) data of every repeat; One-Way ANOVA followed by Tukey’s multiple comparisons test was performed. p-values as indicated on the graph, ns=not significant.

To explore mechanisms behind the reduced metastasis of CKPC mice, we used an *in vivo* transplantation assay to test the metastatic seeding in the peritoneal cavity. This assay also allows us to rule out whether reduced metastasis was just due to the earlier progression to endpoint in CKPC mice. CYRI-B CRISPR (knockout of *cyri-b*, Ex 3) and control KPC-1 cells (Fig S3A) which show similar levels of proliferation (Fig S3B), were injected in the peritoneal cavity of nude mice and metastatic seeding was quantified. Although the pancreas weight to body mass ratio did not change (Fig. 4C), there was a significant reduction in the formation of small metastatic buds on mesentery in the mice injected with CYRI-B CRISPR KPC-1 cells (Fig. 4D,E). No differences in proliferation were observed by Ki67 staining of tumors (Fig. 4F). This mouse model also displays jaundice and ascites fluid, two symptoms which are very common in pancreatic cancer patients. We did not observe any difference in the number of mice presenting with jaundice, however there was a reduction in the number of mice with ascites fluid in mice bearing CYRI-B CRISPR KPC-1 cells (Fig. 4 G,H). Thus, CYRI-B is required for efficient metastatic seeding of KPC cells.

### CYRI-B deletion reduces chemotactic potential

Since we found that CYRI-B can influence the metastatic seeding of KPC cells, we sought to investigate whether CYRI-B can affect their chemotactic potential. Chemotaxis is a major driver of metastasis away from the primary tumour and toward sites rich in attractants, such as blood vessels. It was previously shown that the signalling lipid, lysophosphatidic acid, LPA, which is found in blood serum, is an important chemoattractant driving melanoma and PDAC metastasis (Juin *et al*., 2019; Muinonen-Martin *et al*., 2014). LPA drives chemotaxis of KPC cells and can be sensed by the LPA receptor 1 (LPAR1), present at the plasma membrane of PDAC cells (Juin *et al*., 2019). We generated an independent CKPC cell line, CKPC-1, derived directly from CKPC tumors and rescued with CYRI-B-p17-GFP or GFP (Fig. S4A). To confirm the phenotype, CKPC-1 GFP or rescued cells were seeded on fibronectin-coated glass and stained for ArpC2 to assess the localisation of the Arp2/3 complex at the leading edge (Fig. S4B). CKPC-1 GFP cells presented with more lamellipodia, larger area and increased ArpC2 recruitment to the leading edge in comparison with the rescued cells (Fig. S4B-E) in line with previous results for other cell types (Fort et al., 2018).

Using Insall chemotaxis chambers (Muinonen-Martin *et al*., 2014), we investigated whether CKPC-1 cells can migrate up fetal bovine serum (FBS) gradients, which are a rich source of LPA. Both spider plots and rose plots showing the paths of individual cells and the mean resultant vector of migration, respectively, revealed that CKPC-1 cells have dramatically reduced chemotactic potential towards FBS (Fig. 4I,J). On the contrary, re-expressing GFP-tagged CYRI-B in CKPC-1 cells fully restored chemotaxis and directed migration towards FBS (Fig. 4I,J). CKPC-1 rescued with GFP-tagged CYRI-B were also treated with LPA antagonist KI16425 (Ohta *et al*, 2003) showing that inhibition of LPAR1 and 3 by KI16425 abolished chemotactic steering, consistent with LPA being the major attractant in these conditions (Fig. 4I,J). To further confirm that CYRI-B affects the chemotactic potential of PDAC cells, KPC control and CYRI-B CRISPR cells were also assessed for their chemotactic ability towards serum (10% FBS) using Insall chambers. Deletion of CYRI-B (Ex3 and Ex4) did not change the proliferation rate of cells (Fig. S3), but reduced chemotactic migration in comparison with control cells (Fig. S5). Therefore, CYRI-B is required for chemotactic migration towards serum LPA in PDAC cells.

### CYRI-B localises on macropinocytic cups and vesicles

Having shown that CYRI-B can influence the metastatic seeding of KPC tumors *in vivo* by regulating chemotactic migration, we further probed the role CYRI-B in chemotaxis. We first examined dynamic localisation of CYRI-B, using GFP-labelled CYRI-B (CYRI-B-p17-GFP) and live cell imaging of both COS-7 cells and CKPC cells. Interestingly, CYRI-B was present on internal vesicles and tubules which arise from the vesicles (Fig. 5A and Video S1). The lifetime of vesicular CYRI-B containing structures was around 40s (Fig. 5B), with an average diameter of about 1μm (Fig. 5C), whereas the tubules were up to 17μm in length (Fig. 5D). Additionally, we noticed CYRI-B localising at membrane cups (Fig. 5E and Video S2). CYRI- B positive pseudopods extend nascent cups, fuse together and they slowly move inside the cells with a mean lifetime of about 19s (Fig. 5E). Thus, CYRI-B localised on structures resembling macropinocytic cups, vesicles and associated tubules, similar to what we previously described for CYRI-A (Le *et al*., 2021).

**Figure 5.**
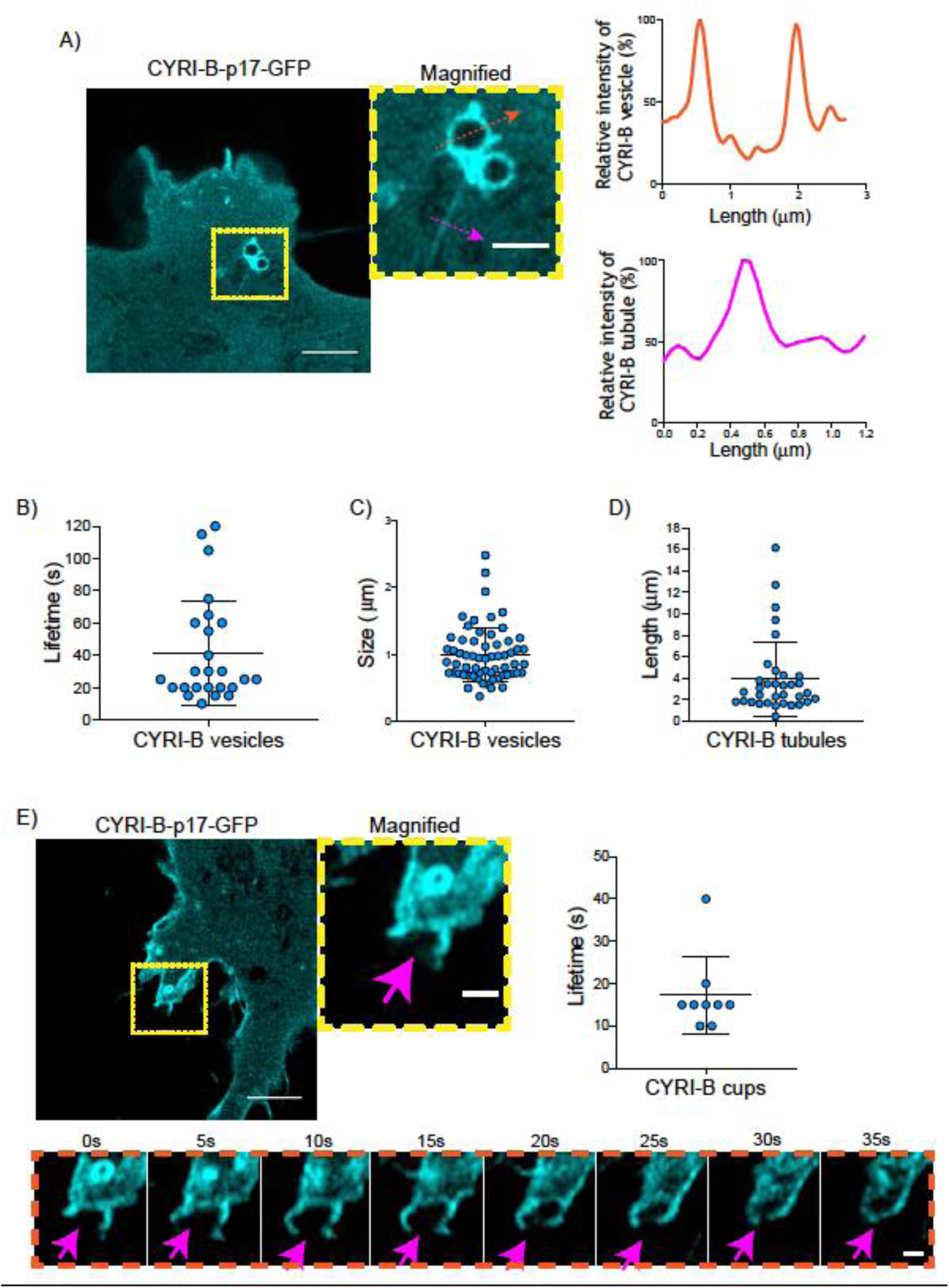
CYRI-B is localised at intracellular vesicles, tubules and membrane cups. A) Still image from live-cell videos of COS-7 CYRI-B KO cells transfected with CYRI-B-p17- GFP (cyan)- see Video 1. Scale bar, 5μm. Yellow box denotes magnified area. Magenta and orange arrows show the quantification area. Scale bar, 1μm. Right panels show the quantifications of the relative intensity of the vesicles/cups and tubules. Image and quantification are representative of n=25 vesicles from a total of 10 cells, over 3 independent biological repeats. B) Scatter plot of the lifetime of vesicles from A). Error bars show the mean ± SD. C) Scatter plot of the size (diameter) of CYRI-B positive vesicles from A). Error bars show the mean ±SD. D) Scatter plot of the length of CYRI-B tubules from A). Error bars show the mean ± SD. E) Still image from live-cell videos of COS-7 CYRI-B KO cells transfected with CYRI-B-p17- GFP (cyan), showing a macropinocytic cup – see Video 2. Scale bar, 5μm. Yellow box denotes magnified area. Magenta arrows show the quantification area. Scale bar, 1μm. Scatter plot on the right panel shows the lifetime of the CYRI-B cups. Error bars show the mean ± SD Orange dotted box shows the montage of the CYRI-B cup over time (s). Scale bar, 1mm. Magenta arrows show the area of interest. Image and quantification are representative of n=9 events from a total of 4 cells.

The CYRI-B positive cups and vesicles were in the size range of macropinosomes (0.2μm- 5μm), rather than other endocytic vesicles and cups, which are typically less than 0.2μm (Canton, 2018). To test the role of CYRI-B in macropinocytosis, we added large molecular weight fluorescently labelled dextran (70kDa) which can only enter by macropinocytosis (Commisso et al., 2013). Live cell imaging of COS-7 cells transfected with CYRI-B-p17-GFP showed that CYRI-B positive pseudopods arising from the membrane enclose dextran, fuse together and internalise (Fig. S6 and Video S3). It was important to test whether this occurred in PDAC cells, since previous work was done in other cell types (Le et al., 2021). CYRI-B stable CKPC-1 PDAC cell line ought to show enhanced macropinocytosis due to the active KRAS (Commisso *et al*., 2013; Kamphorst *et al*., 2015; Palm *et al*, 2017). In agreement with this, PDAC cells showed CYRI-B positive finger-like protrusions extending from the plasma membrane, until they fuse together engulfing extracellular Dextran (Fig. 6A and Video S4). The CYRI-B positive macropinosomes were then internalised and travelled inside the cells until they disappeared. Quantification of the lifetime of CYRI-B macropinosomes showed very similar results between the two cell lines (Fig. 6A). Thus CYRI-B localises on macropinosomes in PDAC cells, suggesting a possible mechanism for how CYRI-B loss could affect tumour progression.

**Figure 6.**
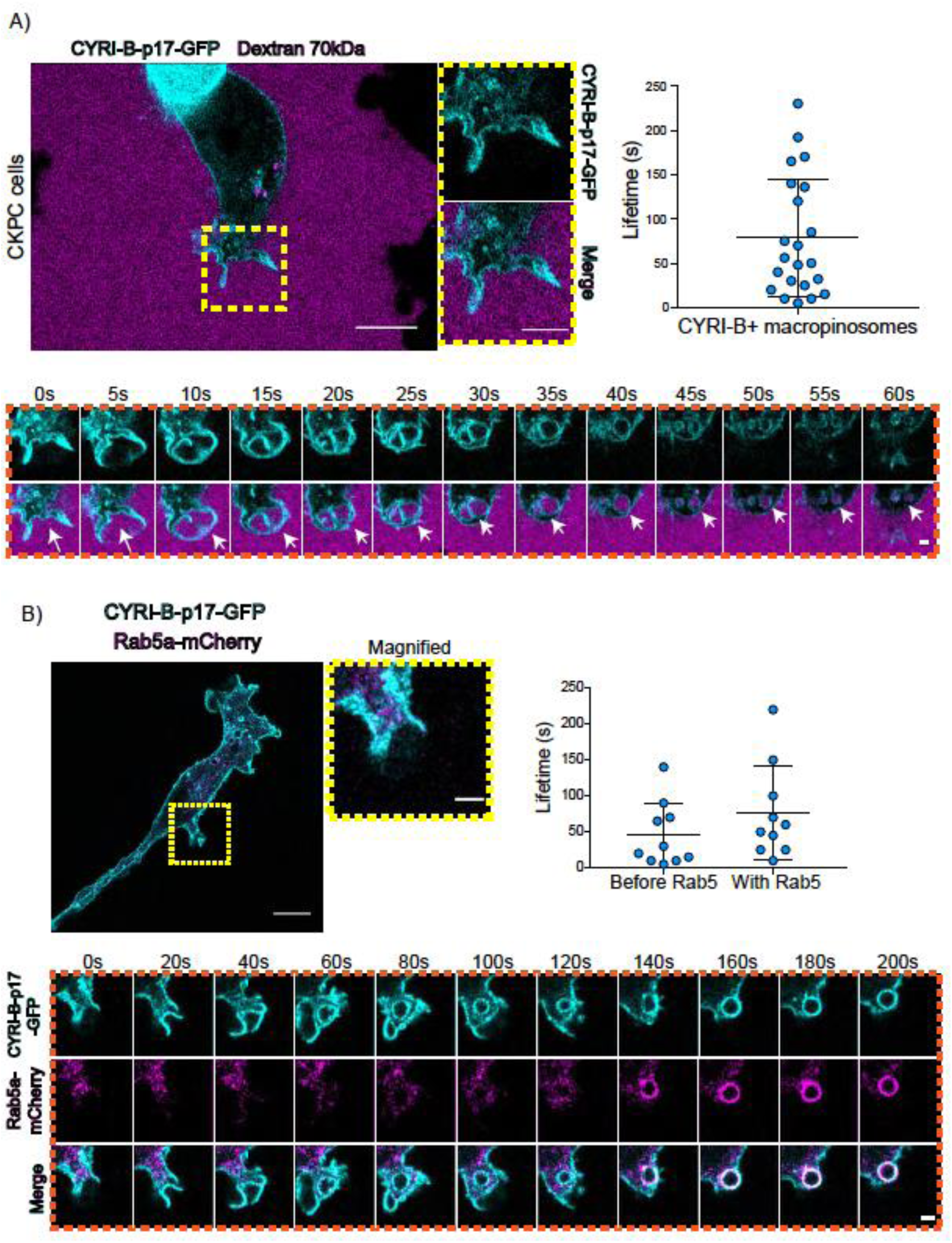
CYRI-B is recruited to macropinocytic cups and precedes Rab5 recruitment. A) Still image from live-cell imaging of CKPC-CYRI-B-GFP stable cell lines (cyan)- see Video 3. 70kDa Dextran was added to the medium to visualise macropinocytic events (magenta). Scale bar, 10μm. Yellow box shows the magnified area of interest, showing the macropinocytic cups. Scale bar, 5μm. Scatter plot represents the lifetime of CYRI-B+ macropinosomes once internalised. Mean ± SD. Orange box shows a representative montage of CYRI-B internalisation via macropinocytosis. Scale bar, 1mm. White arrows show CYRI-B localisation at the cups and the macropinosomes once internalised. n=21 events from a total of 6 cells. B) Still image from live-cell imaging of COS-7 CYRI-B KO cells transfected with CYRI-B-p17- GFP (cyan) and mRFP-Rab5 (magenta)- See Video 5. Scale bar, 10μm. Yellow box show the magnified area of interest, showing the macropinocytic cups. Scale bar, 5μm. Orange boxes show a representative montage of CYRI-B internalisation and the recruitment of Rab5 at the nascent macropinosomes. Scale bar, 5mm. Scatter plots represent the lifetime of CYRI-B+ macropinosomes once internalised before and after Rab5 recruitment. Error bars show the mean ± SD; n=10 events from a total of 6 cells.

One of the first proteins to be recruited to macropinosomes once they internalise is Rab5, which is present on vesicles that move from the periphery of the cells towards the perinuclear region (Bucci *et al*, 1994; Buckley & King, 2017; de Hoop *et al*, 1994). We found previously that CYRI-A showed a transient recruitment to macropinocytic cups and was largely absent from macropinosomes that had internalised, as marked by recruitment of the early endosome component Rab5 (Le et al., 2022). Therefore, we examined the localisation of CYRI-B relative to the early endosome component Rab5. Live-cell imaging of COS-7 cells transfected with both CYRI-B-p17-GFP and Rab5-mcherry showed that Rab5 is recruited after CYRI-B macropinosome internalisation. First CYRI-B positive pseudopods extend and fuse together to form the nascent macropinosome which is then internalised (Fig. 6B and Video S5). After ∼50s of internalisation, Rab5 is recruited to the macropinosomes (Fig. 6B and Video S5) suggesting that CYRI-B is present prior to and also during early macropinosome formation as marked with Rab5.

### LPAR1 internalises via CYRI-B positive macropinosomes

An important but often overlooked role of macropinocytosis is the maintenance of cell surface receptors (Buckley & King, 2017). Chemotaxis toward LPA requires the fine coordination of multiple proteins at the cell leading edge in order to sense LPA, internalise the LPAR1 receptor and recycle it back to the plasma membrane (Juin *et al*., 2019; Muinonen-Martin *et al*., 2014). Having optimal amounts and dynamics of LPAR receptors at the leading edge is critical for a coordinated movement of cells toward the chemoattractant (Juin *et al*., 2019). Therefore, we hypothesized that CYRI-B might influence the internalisation of the LPA receptor at the leading edge. 70kDa TRITC dextran was added to the medium and cells were imaged using live time lapse microscopy to visualise the macropinosomes in COS-7 cells transfected with LPAR-1 GFP. We observed that LPAR1-positive vesicles incorporated dextran and after some time they disappeared, with a mean lifetime of ∼58s (Fig. 7A). Thus, LPAR1 is taken up from the cell surface by macropinocytosis.

**Figure 7:**
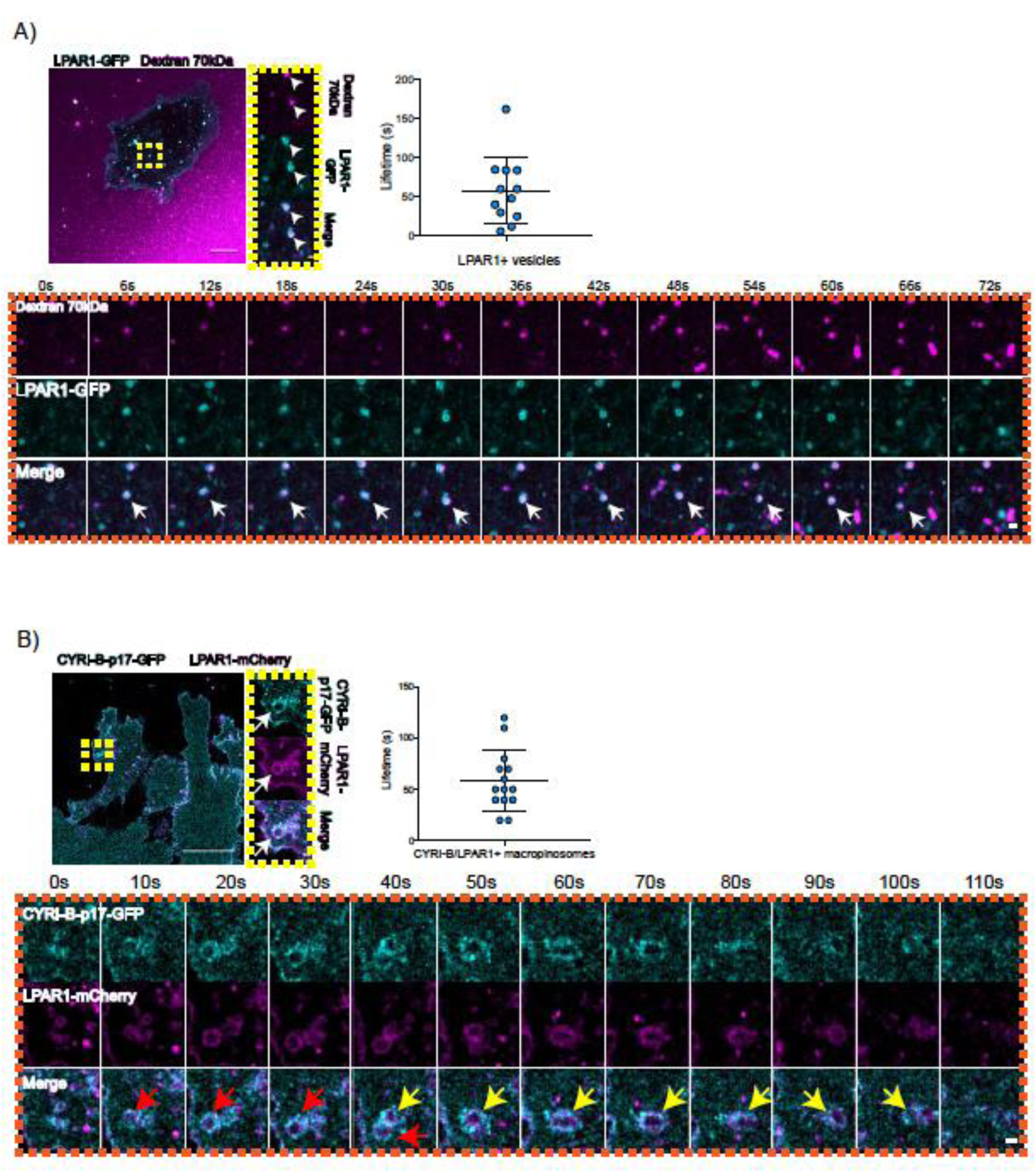
LPAR is internalised via CYRI-B positive macropinocytosis. A) Still images from live-cell imaging of COS-7 cells transfected with LPAR1-GFP (cyan)- see Video 6. 70kDa Dextran was added to the medium to visualise the macropinosomes (magenta). Scale bar, 10μm. Yellow box shows the magnified area of interest, showing the LPAR1+ macropinocytic vesicles/cups. White arrows denote structures of interest. Scale bar, 1μm. Scatter plot represents the lifetime of LPAR1+ vesicles once internalised. Mean ± SD. Orange box shows a representative montage of LPAR1 internalisation via macropinocytosis. Scale bar, 1mm. White arrows show the vesicle of interest. n=12 events from a total of 3 cells. B) Still image from live-cell imaging of CKPC-1 cells transfected with CYRI-B-p17-GFP (cyan) and LPAR1-mCherry (magenta)- see Video 7. Scale bar, 20μm. Yellow box shows the magnified area of interest, showing the LPAR1 co-localisation with CYRI-B+ macropinosomes. White arrows show the vesicle of interest. Scale bar, 1µm. Scatter plot represents the lifetime of LPAR1 and CYRI-B vesicles once internalised. Mean ±SD. Orange box shows a representative montage of LPAR1 and CYRI-B internalisation. Red and yellow arrows show the vesicles of interest. n=14 events from a total of 4 cells.

To ask whether the LPAR-1-positive macropinocytic structures also contained CYRI-B, we transfected CYRI-B-GFP stable CKPC-1 cells with LPAR1-mCherry and performed live-cell imaging. Indeed, upon CYRI-B internalisation, LPAR1 is also internalised with a lifetime of ∼58s, consistent with LPAR-1 trafficking via CYRI-B positive macropinocytic events in PDAC cells (Fig. 7B). Thus, LPAR1 is at least partially internalised via CYRI-B mediated macropinocytosis.

### CYRI-B controls chemotactic migration via macropinocytic LPAR-1 internalisation and membrane localisation

Having found that CYRI-B co-localises with LPAR1 and deletion of CYRI-B affects the chemotactic ability of PDAC cells to migrate in vitro and in vivo, we investigated whether CYRI-B could influence the trafficking of LPAR-1. Initial work showed that CKPC-1 cells expressing either GFP or CYRI-B-p17-GFP following stable transfection did not alter mRNA levels of LPAR1 or LPAR3 (Fig. S7A) suggesting that changes in the chemotactic ability of the cells is not likely due to alterations in the expression level of the LPARs. Since CYRI-B alters the shape of CKPC cells we also checked whether the localisation of LPAR is changed. CKPC-1 cells (with stable expression of GFP or CYRI-B-p17-GFP) were transfected with HA- LPAR1 and fixed for immunofluorescence. Cells displayed localisation of HA-LPAR1 at the plasma membrane and internal vesicles, as expected from previous reports (Juin *et al*., 2019). CYRI-B knockout cells (expressing GFP-only) showed high levels of leading-edge membrane localisation of LPAR1 in comparison with the CYRI-B-p17-GFP rescued cells (Fig. S7B and C).

The combined evidence of the role of CYRI-B on macropinocytic uptake, the co-internalisation of CYRI-B with LPAR1 as well as the increase in peripherally accumulated LPAR1, led us to ask whether CYRI-B can affect the internalisation LPAR1. We performed an image-based internalisation assay, using CKPC-1 stable GFP and CYRI-B-p17-GFP cells that were transfected with LPAR-1-mCherry and serum-starved overnight. Cells were stimulated with 10% FBS for 15min and fixed to measure the internalisation of LPAR1. Previous reports suggested that stimulation of cells with serum should cause an increase in the internalisation of GPCRs including LPAR1 (Juin *et al*., 2019; Kang *et al*, 2014). CYRI-B rescued cells showed a serum-stimulated enhancement of LPAR1 internalisation, while CYRI-B depleted cells showed minimal stimulation of LPAR1 uptake (Fig. 8A,B). While LPAR1 may be internalised by multiple endocytic pathways, the dependence on CYRI-B suggests that LPAR- 1 is a cargo of CYRI-B dependent macropinocytosis, regulating surface levels and chemotactic migration.

**Figure 8:**
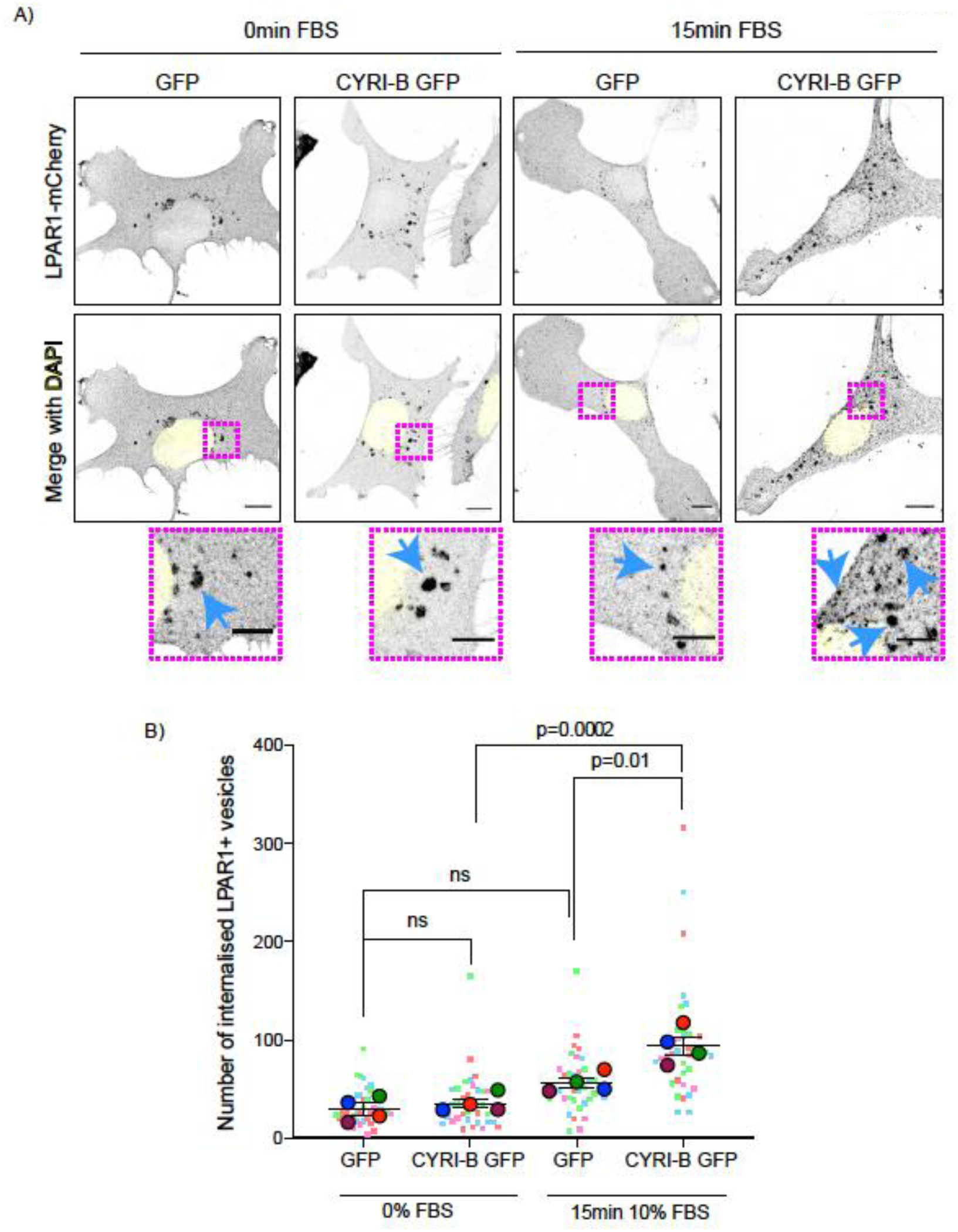
Loss of CYRI-B reduces LPAR1 internalisation upon serum stimulation. A) Immunofluorescence images of CKPC-1 stable cells transfected with GFP or CYRI-B-p17- GFP. Cells were transfected with LPAR1-mCherry and seeded on fibronectin coated coverslips. Cells were starved overnight and the next day 10% FBS was used to stimulate the uptake of LPAR1. Vesicles (marked by LPAR1-mCherry) are shown as black dots, DAPI (yellow) was used to visualise the nuclei. Scale bars 10µm. Magenta dotted boxes show the magnified area of interest and cyan arrows show the internalised vesicles. Scale bars 5µm. B) The graph shows the quantification of the number of LPAR1+ vesicles in each condition. Scatter plot is presented as super plots and every independent biological repeat is coloured differently. Mean ± S.E.M; One-Way ANOVA followed by Tukey’s multiple comparisons test was performed, n=4 (from a total of ≥35 cells for each condition). p-value as indicated, ns = not significant

## Discussion

We have revealed an important role of CYRI-B in PDAC development, progression and metastasis using the KPC mouse model and cells derived from the tumours. Our previous cell biology studies highlighted a role for CYRI-B as a buffer of RAC1-mediated actin assembly in lamellipodia and macropinocytic cups (Fort *et al*., 2018; Le *et al*., 2021), but did not address the potential role that CYRI-B could play in tumourigenesis and metastasis, given its central role as a regulator of motility. We noticed that CYRI-B was highly expressed in human pancreatic cancers and correlated with poorer survival (Nikolaou & Machesky, 2020). Increased expression of CYRI-B in mice with PDAC suggested that it would be worth exploring possible functional consequences of CYRI-B loss in the KPC mouse model.

Deletion of *cyri-b* in the pancreas, in concert with expression of KRAS ^G12D^ and p53 ^R172H^ led to acceleration of PanIN formation and an increase in the area of pancreas showing lesions with high phospho-ERK and Phospho-JNK, two crucial downstream targets of KRAS and RAC1 that drive proliferation and expansion. Other recent studies implicated CYRI-B (Fam49B) autoantibodies as a potential biomarker for early stage breast cancer (Luo *et al*, 2022), a gene found in patient serum on extrachromosomal circular DNA overexpressed in lung adenocarcinoma (Xu *et al*, 2022b) and a potential saliva marker of oral cancer (Kawahara *et al*, 2016). CYRI-B was also highlighted as a target of the zinc finger RNA-binding protein Zfrbp, leading to accelerated tumour development when overexpressed in colorectal and liver cancers (Long *et al*, 2019). Further investigations are needed to determine the nature of CYRI- B as a potential biomarker of early cancer, but multiple studies suggest that it may be enriched in extracellular vesicles associated with cancer and other diseases e.g. (Peng *et al*, 2019). Taken together, CYRI-B may have potential as a biomarker and driver of early cancer and play a role in the progression or conversion from precancer to cancerous lesions.

Another possible mechanism by which loss of CYRI-B could enhance early cancer progression would be via its role in maintenance of epithelial apico-basolateral polarity (Fort *et al*., 2018). We previously found that loss of CYRI-B in MDCK cell spheroids disrupted lumen formation in a similar way to hyperactivation of RAC1. Likewise, RAC1 and PI3-kinase are important for apicobasal polarity in the pancreas (Lof-Ohlin *et al*, 2017) and RAC1 has a known role in acinar to ductal metaplasia and in polarity and cell identity during early PDAC progression (Heid *et al*., 2011). PI3-kinase plays an important role in polarity maintenance and CYRI-B has been implicated in PI3-kinase signaling in gallbladder cancer cells (Zhang *et al*, 2020). Loss of CYRI-B could therefore lead to hyperactivation or inappropriate spatial control of RAC1 activation causing a loss of normal cell polarity and therefore enhancing preneoplasia and cancer progression. Polarity maintenance could be disrupted by a lack of proper control of Scar/WAVE complex localisation leading to aberrant actin regulation (Fort *et al*., 2018), or due to aberrant membrane trafficking of receptors such as integrins (Le *et al*., 2021) or LPAR1 (this study).

Involvement of CYRI-B in chemotactic migration suggested a mechanism for the reduced metastasis that we observed in both the KPC model and the intraperitoneal transplant model of PDAC metastasis. We previously found that loss of N-WASP in the KPC model caused a reduction in metastasis and that the role of N-WASP in recycling the LPA receptor LPAR1 was crucial in mediating this phenotype (Juin *et al*., 2019). While N-WASP localises to SNX9- positive membrane tubules that are involved in trafficking of LPAR1 back to the plasma membrane after internalisation, we find that CYRI-B is required for efficient internalisation of LPAR1 after stimulation. Thus, CYRI-B and N-WASP control two different aspects of a similar pathway, whereby LPAR1 is stimulated, internalised and then either sorted into tubules for rapid recycling or targeted toward lysosomes for slow recycling or degradation.

Interestingly, a major mechanism for internalisation of LPAR1 appears to be via macropinocytosis, as we observed LPAR1 on the membrane surface of both nascent and internalised macropinocytic structures co-localising with CYRI-B. We also found a significant retardation of LPAR1 internalisation in CYRI-B depleted cells, indicating that LPAR1 is significantly internalised via CYRI-B dependent macropinocytosis in PDAC cells. This suggests that macropinocytosis, which has recently attracted substantial interest as a regulator of nutrient uptake by PDAC cells (Canton, 2018; Commisso *et al*., 2013; Michalopoulou *et al*, 2020; Puccini *et al*., 2022; Yao *et al*., 2019) is also a key mechanism by which cells control surface receptor trafficking. Although involvement of macropinocytosis in receptor trafficking has been previously observed (reviewed in (Stow *et al*., 2020)), studies are primarily in immune cells, which perform high levels of constitutive uptake. Macropinocytosis as a way for cancer cells to control signalling and adhesion is perhaps under-appreciated and warrants further study in this capacity.

## Materials and Methods

### Mammalian cell culture

Mammalian cells were cultured with Dulbecco’s Modified Eagle’s Medium (DMEM) (#21969-035; Gibco) growth medium supplemented with 10% FBS (#10270-106; Gibco) and 2mM L-Glutamine (#25030-032; Gibco). The cell lines were split roughly every two days and maintained at 37°C humidified incubator and perfused with 5% CO_2_. For the proliferation assays, approximately 10^4^ KPC CRISPR or control cells were seeded on 6-well plates and were manually counted every day for 4 days.

### Cell transfection

About 1x10^6^ COS-7 cells were transfected with Lipofectamine 2000 (#11668019, Invitrogen) according to manufacturer’s instructions. For KPC-1 and CKPC-1 cell lines, the AMAXA-V kit (VCA-1003, Lonza) was used according to the manufacturer’s protocol. About 2x10^6^ cells were electroporated using P-031 program from the AMAXA electroporator. The transfected cells were left overnight in full media in a humidified incubator at 37°C supplied with 5% CO_2_.

### sgRNAs and KPC CRISPR cell line generation

sgRNAs for CRISPR were designed using the Zhang laboratory website (https://zlab.bio/guide-design-resources). Mouse Cyri-b exon3 (CACCGGGTGCAGTCGTGCCACTAGT) and exon4 (CACCGCGAGTATGGCGTACTAGTCA) were used for CrispR cell line generation and transfected into lentiCRISPRv1-puro.

To generate CYRI-B knockout stable cell lines, CRISPR-Cas9 genome editing technology was performed, using the calcium phosphate transfection kit (#K2780-01, Invitrogen). To generate the virus which infected the recipient cells (KPC or CKPC cells) the HEK293T cell line was used. First, about 2x10^6^ HEK293T cells per 10cm dish were seeded and let overnight to grow. Next day the transfection master mix which contained 10µg of CRISPR construct containing sgRNA targeting the gene of interest (or empty lentiCRISPRv1-puro, #49535, Addgene), 7.5µg of pSPAX2 (#12260, Addgene), and 4µg of pVSVG packaging plasmid (#8454, Addgene) was prepared according to manufacturer’s instructions. The following day the medium was removed and replaced with the same medium composition (DMEM) with 20% FBS, for virus production. The cells were left overnight and in the meantime recipient cells were prepared for virus infection by seeding 1x10^6^ cells per plate. The next day the medium from the HEK293T cells was removed and mixed with 2.5μl hexadimethrine bromide (10mg/mL) (#H9268, Sigma), filtered using a 0.45μm pore membrane to remove any cell debris. The medium was then added to the recipient cells and left overnight. The next day the same procedure was repeated to achieve better infection with the virus. Transduced cells were selected using puromycin (2μg/ml) (#ant-pr-1; InvivoGen).

### CKPC cell lines generation

CKPC cell lines (CKPC-1 and 2) were first generated by taking about 1/3 of the tumors from two different end-point mice. The tumors were washed three times with 5% penicillin/streptomycin (#15140122; Life Technologies) in PBS and cut into small pieces (<3mm). The tumor pieces were then washed with PBS, centrifuged for 5min at 1200 rpm and transferred to 10cm plates using full DMEM media supplemented with primocin (1:1000). The plates were left overnight in humidified incubator at 37°C supplied with 5% CO_2_ until confluent. After about 5-7 passages, cells were checked for Pdx-1, p53 and CYRI-B protein staining.

For CYRI-B rescued stable cell line creation, cells were transfected with CYRI-B-p17-GFP along with a puromicin resistant plasmid using AMAXA-V kit as previously described. For control purposes the same CKPC cell line was also transfected with the empty GFP backbone. Cells were selected using 1mg/ml puromicin and FACs sorted. Low-medium intensity GFP positive cells were selected to ensure that CYRI-B is not overexpressed. Cells were checked for CYRI-B expression and kept for maximum of 3-4 passages.

### Chemotaxis assay

Chemotaxis assay was performed as previously described in (Muinonen-Martin et al, 2010). About 2x10^5^ cells were seeded on coverslips. Following attachment, the medium was replaced with SFM DMEM to starve the cells and left overnight. Next day the “Insall” chemotaxis chambers were prepared. In the middle chamber, serum starvation medium was added. The coverslip was then carefully placed cell-side down onto the chamber. To create a chemoattractant gradient, full DMEM medium with 10% FBS was added on the sides of the Insall chambers (about 120μl). The bridges containing the cells where then visualised every 15min using a Nikon long term time-lapse microscope for 48h. For LPAR1/3 inhibitor treatment with KI16425 antagonist (Cayman chemicals, #10012659), cells were incubated for 1h in serum free medium in 1:1000 dilution prior preparation and assembly of the “Insall” chambers. DMSO (#15572393, Fisher Chemical) was used as a vehicle control. The cells where manually tracked using the mTrackJ plugin of Fiji software. From each condition, at least two random bridges where selected and at least 25 cells from each bridge where manually tracked. Only cells present at the first frame of the video were counted in the tracking, whereas when the cells were moving outside the bridge the tracking was stopped. The Excel spreadsheets with all the cell tracks from each bridge, were extracted in order to create rose- plots, individual cell-track graphs and cosθ data using an algorithm in R software which was previously designed and published in our lab (Fort *et al*., 2018).

### Western blotting

Protein extraction from cultured cells was performed using ice-cold RIPA (150 mM NaCl, 10 mM Tris-HCl pH 7.5, 1 mM EDTA, 1 % Triton X-100, 0.1 % SDS buffer) supplemented with 1x phosphatase and 1x protease inhibitors (#78427, #78438; Thermo Fisher Scientific). Cell lysates were centrifuged at 13000rpm at 4°C for 10min and the supernatant was collected. Protein was quantified using the Precision Red (#ADV02; Cytoskeleton) advanced protein assay and 10-20μg was used. The lysates were mixed with 1x NuPAGE LDS sample buffer (#NP0007, Invitrogen) and 1x NuPAGE reducing agent (#NP0004, Invitrogen), boiled for 5min at 100°C and loaded on Novex 4-12% Bis-Tris acrylamide pre-cast gels (#NP0321; Thermo Fisher Scientific) at 170V for about 1h. The proteins were transferred onto a 0.45μm nitrocellulose blotting membrane (#10600002; GE Healthcare) using wet electrotransfer for 1h at 110V. The membranes were blocked with 5% (w/v) BSA diluted in 1x TBS-T (10mM Tris pH 8.0, 150mM NaCl, 0.5 % Tween-20) for 30min at room temperature on a shaker. Primary antibodies were incubated in buffer with 5% (w/v) BSA and 1x TBS-T overnight at 4°C on a roller. Membranes were washed 3 times and incubated with Alexa-Fluor conjugated secondary antibodies (#A21206 and #A10038, Thermo Fisher Scientific) diluted in 5% (w/v) BSA and 1x TBS-T, for 1h at room temperature on a roller. The membranes were then washed 3 times and visualised using the Li-Cor Odyssey CLx scanner with the auto intensity scanning mode.

### Immunofluorescence assay

CKPC cells were plated onto sterile 13 mm glass coverslips that had been previously coated with 1mg/ml fibronectin (#F1141; Sigma-Aldrich). Coverslips were fixed using 4% PFA (#15710, Electron Microscopy Sciences) in PBS for 10min at room temperature. The coverslips were then washed 3 times with PBS and permeabilised with permeabilisation buffer for 5min at room temperature. The coverslips were washed again 3 times with PBS and blocked with blocking buffer for about 30min at room temperature. Primary antibodies were diluted in blocking buffer in the appropriate dilution and incubated for 1h at room temperature. Coverslips were washed 3 times with blocking buffer and secondary antibodies were then added in the appropriate dilution in blocking buffer and incubated for 1h at room temperature. Finally, the coverslips were washed 3 times with PBS and mounted on microscopy slides using FluoromountG solution containing DAPI (Southern Biotech; 0100-01). Slides were imaged using a Zeiss 880 LSM with Airyscan microscope.

### Live-cell imaging

For live cell imaging cells were seeded on glass bottom plates which were previously coated with either 1mg/ml fibronectin (CKPC cells) or 10 μg/ml laminin (COS-7 cells, #L2020, Sigma). The cells were imaged using a Zeiss 880 LSM with Airyscan microscope which has a 37°C humidified incubator and perfused with 5% CO_2_. Depending on the experiment either still images or videos were taken as mentioned in the results chapters.

### Image-based LPAR1 internalisation assay

CKPC-1 stable cell lines were first transfected with LPAR1-mCherry construct. The cells were then seeded on fibronectin coated coverslips and once adhered the media was replaced with serum-free DMEM overnight. The following day 10% FBS was added to the pre-existing medium and incubated for 15min for the internalisation to occur. The media was aspirated and the immediately fixed using 4% PFA. The cells were then mounted on microscopy slides and visualised using a Zeiss 880 LSM with Airyscan microscope (x63 objective). Images were analysed by thresholding the LPAR1-mCherry channel and analysing the objectives which were 0.1 or above in Fiji software.

### Reverse transcription quantitative Polymerase Chain Reaction (RT-qPCR)

RNA was first isolated from the cells using the RNeasy Mini Kit (#74104, Qiagen) according to the manufacturer’s instructions. The cDNA was then synthesised using the SuperScript III Reverse Transcriptase protocol and measured using NanoDrop2000c. 1μg of cDNA was mixed along with a primer master mix previously made. The qPCR was performed according to manufacturer’s instructions using and the DyNAmo HS SYBR Green qPCR kit (#F410L; Thermo Fisher Scientific) reaction was set up as follows using the C1000 Thermal Cycler (CFX96 Real time system, BioRad): 3 min at 95°C, 20 s at 95°C, 30 s at 57°C, 30 s at 72°C, repeat steps 2-4 for 40 cycles and 5 min at 72°C. Each condition had three technical replicates and GAPDH was used as a housekeeping gene (Fw – CATGGCCTACATGGCCTCCA, Rv- TGGGATAGGGCCTCTCTTGC, ThermoFisher). The mRNA fold change was calculated using the ΔΔC_t_ method. The LPAR1 and LPAR3 Quantitec primers were purchased from Qiagen (#QT00264320, #QT00107709).

### Mouse model

The mice were maintained by the Biological Services Unit staff according to the UK home office regulations and instructions. The experiments were approved by the local Animal Welfare and Ethical Review Body (AWERB) of the University of Glasgow and performed under UK Home office licence PE494BE48 to LMM. Data for cohorts is included in the Supplementary Spreadsheet Table S1. The genotyping was performed by ear notch and the samples were sent to TransnetYX®. To generate the *Cyri-b* floxed (Cyri-b^fl/fl^) mouse, frozen sperm was obtained from the Canadian Mouse Mutant Repository (Fam49b_tm1c_C08). The mouse strain was generated by IVF (Takeo & Nakagata, 2011, 2015) using C57BL/6J mice as embryo donors, and the resulting 2-cell embryos transferred to pseudopregnant recipients using standard protocols. The CKPC mouse model was generated by crossing *LSL-KRAS^G12D^, LSL- p53^R172H^, Pdx1::CRE* (KPC) mice (Hingorani *et al*, 2003) with *cyri-b* floxed (*cyri-*b^fl/fl^) mice. Mice that died from causes other than pancreatic cancer were removed from the study.

### In vivo transplantation assay

For the *in vivo* transplantation assay KPC-1 CYRI-B CRISPR cell lines were used. The cells were grown in full media as normal until 24 hours before transplant, when fresh media was added without any antibiotics. About 2x10^6^ cells per mouse were used for the experiment. Cells were injected into the intraperitoneal cavity of 10-week old female CD-1 mice (Charles Rivers). Once injected, the mice were monitored every day and at day 14 mice were sacrificed. The weight of the mice and pancreata were taken at end-point.

### Immunohistochemistry, In Situ Hybridization detection (RNAScope) assays and quantification

Tissues were fixed in 10% formalin and next day transferred to 70% ethanol. The tissues were then embedded into paraffin blocks. For Immunohistochemistry, the staining was performed on 4µm sections which had previously been ovened at 60⁰C for 2 hours, using standard protocols. The detection for Mm-*cyri-b* mRNA was performed using RNAScope 2.5 LS (Brown) detection kit (#322100; Advanced Cell Diagnostics, Hayward, CA) on a Leica Bond Rx autostainer strictly according to the manufacturer’s instructions. Slides were imaged using the Leica SCN 400f scanner.

To quantify the histology slides, the HALO software was used. About 8 different areas (>350000µm^2^ each area) within the pancreatic tumours were used to quantify the different stains. Necrotic areas were quantified manually from the whole pancreatic tumour using H&E staining. Areas with fragmented nuclei were considered as positive for necrosis. For tissues from 15-week old mice, both neoplastic lesions and tumour areas were quantified (pJNK, pERK, BrdU stains). For the RNAScope experiments the algorithm was set up to recognise the individual dots. PanIN lesion quantification was performed manually according to the following website: https://pathology.jhu.edu/pancreas/medical-professionals/duct-lesions using H&E slides.

### Statistical analysis

Data and statistics were analysed using the Prism software. All of the cell biology experiments, unless otherwise stated in the figure legends, were performed three times on separate occasions with separate cell passages. For all the mouse experiments and histology quantifications for end-point mice between KPC and CKPC mouse models, similar ages of mice were chosen. Unless otherwise stated, all the cell biology experiments were plotted as superplots (Lord *et al*, 2020) and each biological replicate was coloured differently. The quantified numbers of individual cells from the repeats are shown as individual points in the background. smaller shapes and coloured depending on the repeat. Generated graphs show either S.E.M. or S.D. depending on the experiment. n.s.=not significant/p>0.05, *p<0.05, **p≤0.01, ***p≤0.001, ****p≤0.0001, unless the exact p-value is shown on the graph.

## Data Availability

The original data for this paper has been deposited to the University of Glasgow’s Enlighten Database with the DOI http://dx.doi.org/10.5525/gla.researchdata.1371

## Acknowledgements

We thank Cancer Research UK for core funding (A17196 and A31287) and funding to L.M.M. (A24452) and UKRI EPSRC grant (EP/T002123/1) to L.M.M.

## Appendix- Key resources tables

### Antibodies and Reagents for Western blotting or Immunofluorescence or live-cell imaging

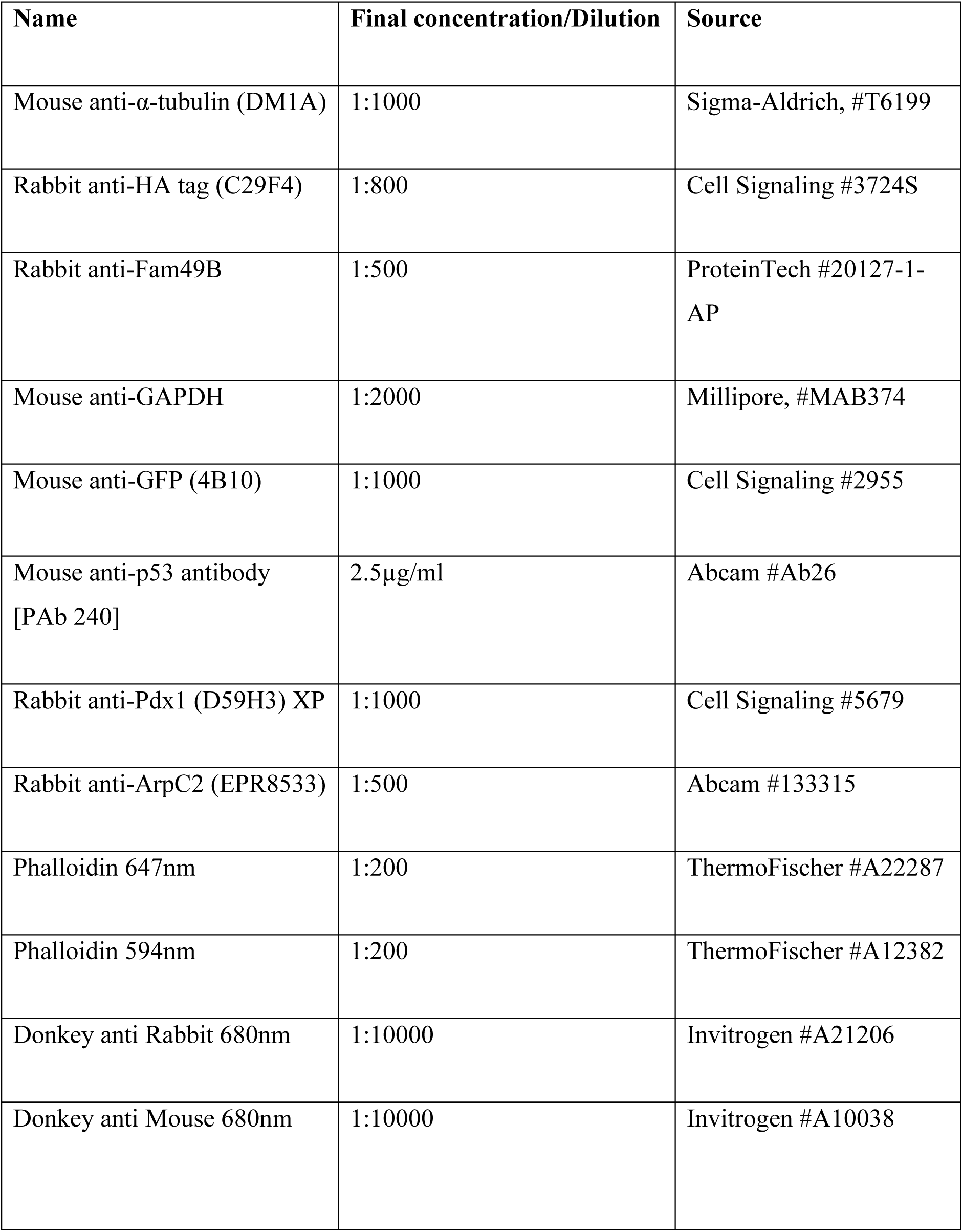
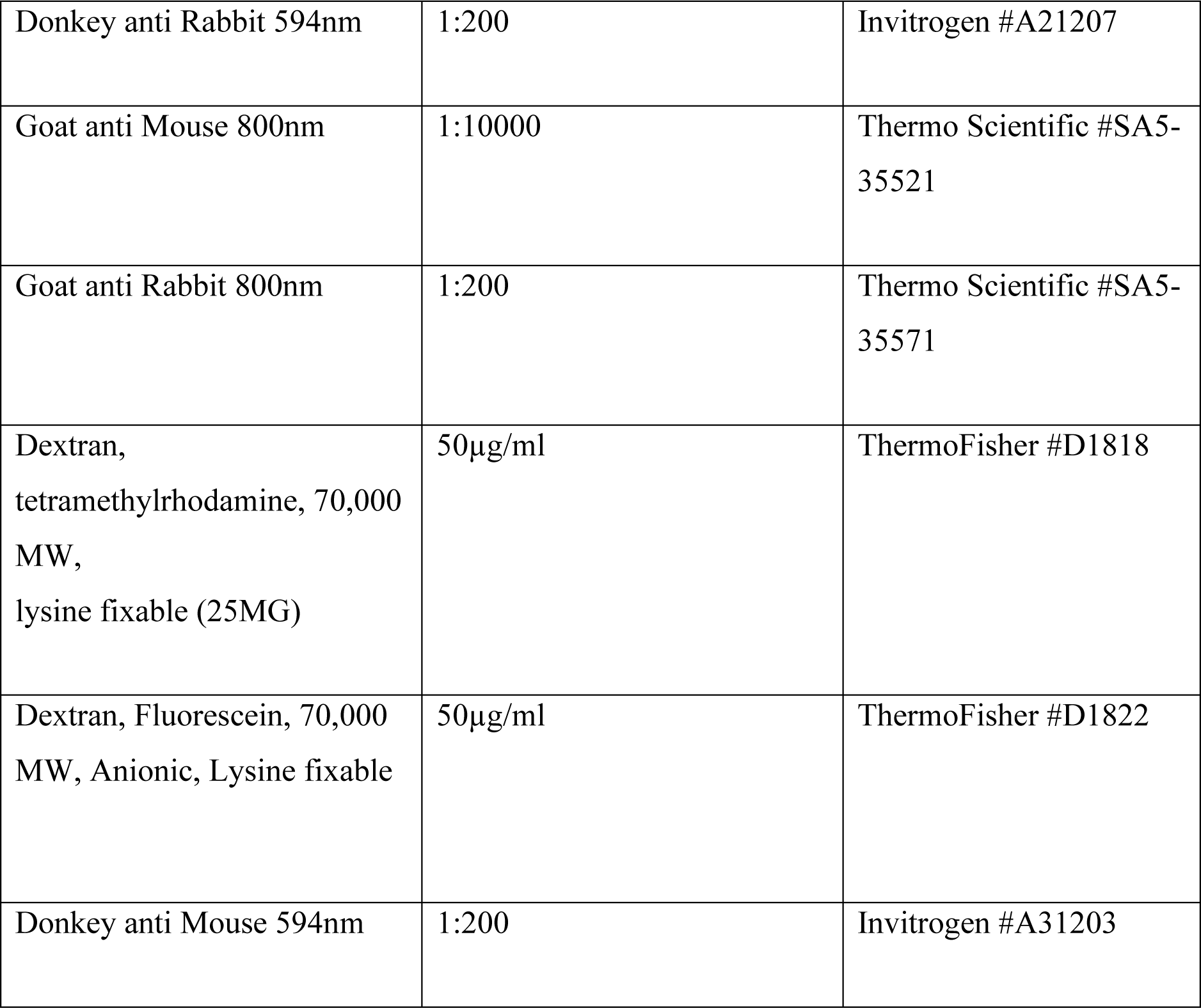

| **Name** | **Final concentration/Dilution** | **Source** |
| --- | --- | --- |
| Mouse anti-α-tubulin (DM1A) | 1:1000 | Sigma-Aldrich, #T6199 |
| Rabbit anti-HA tag (C29F4) | 1:800 | Cell Signaling #3724S |
| Rabbit anti-Fam49B | 1:500 | ProteinTech #20127-1-AP |
| Mouse anti-GAPDH | 1:2000 | Millipore, #MAB374 |
| Mouse anti-GFP (4B10) | 1:1000 | Cell Signaling #2955 |
| Mouse anti-p53 antibody [PAb 240] | 2.5μg/ml | Abcam #Ab26 |
| Rabbit anti-Pdx1 (D59H3) XP | 1:1000 | Cell Signaling #5679 |
| Rabbit anti-ArpC2 (EPR8533) | 1:500 | Abcam #133315 |
| Phalloidin 647nm | 1:200 | ThermoFischer #A22287 |
| Phalloidin 594nm | 1:200 | ThermoFischer #A12382 |
| Donkey anti Rabbit 680nm | 1:10000 | Invitrogen #A21206 |
| Donkey anti Mouse 680nm | 1:10000 | Invitrogen #A10038 |
| Donkey anti Rabbit 594nm | 1:200 | Invitrogen #A21207 |
| Goat anti Mouse 800nm | 1:10000 | Thermo Scientific #SA5-35521 |
| Goat anti Rabbit 800nm | 1:200 | Thermo Scientific #SA5-35571 |
| Dextran, tetramethylrhodamine, 70,000 MW, lysine fixable (25MG) | 50µg/ml | ThermoFisher #D1818 |
| Dextran, Fluorescein, 70,000 MW, Anionic, Lysine fixable | 50µg/ml | ThermoFisher #D1822 |
| Donkey anti Mouse 594nm | 1:200 | Invitrogen #A31203 |

### Antibodies and Reagents for IHC

| Name | Dilution | Source |
| --- | --- | --- |
| BrdU (B44) | 1:250 | BD Biosciences #347580 |
| Caspase 3 (ASP-175) | 1:500 | Cell Signaling #9661 |
| Ki67 (D3B5) | 1:1000 | Cell Signaling #12202 |
| p44/42 MAPK (ERK1/2) | 1:400 | Cell Signaling #9101 |
| p53 (CM5) | 1:750 | Leica #NCL-L-p53CM5p |
| PDX1 | 1:400 | Abcam #ab47267 |
| pSapk/Jnk Thr183/Thr185<br>(81E11) | 1:20 | Cell Signaling #4668 |
| Flex Wash buffer | N/A | Agilent #K8007 |
| High pH Target Retrieval<br>Solution (TRS) | N/A | Agilent #K8004 |
| Liquid DAB | N/A | Agilent #K3468 |
| Low pH Target Retrieval<br>Solution (TRS) | N/A | Agilent #K8005 |
| Mouse EnVision | N/A | Agilent #4001 |
| Peroxidase block | N/A | Agilent #S2023 |
| Rabbit EnVision | N/A | Agilent #K4003 |
| Rabbit signal boost (HRP) | N/A | Cell Signaling #8114 |
| Bond Wash | N/A | Leica #AR9590 |
| Enzyme pre-treatment kit | N/A | Leica #AR9551 |
| Epitope Retrieval Solution 1<br>(ER1) | N/A | Leica #AR9661 |
| Epitope Retrieval Solution 2<br>(ER2) | N/A | Leica #AR9640 |
| Intense R Kit | N/A | Leica DS9263 |
| Mouse Ig Blocking reagent | N/A | Vector labs #MKB-2213 |
| Rat ImmPRESS kit | N/A | Vector labs #MP-7404 |

### Cell lines and experimental model cell lines

| Name | Source |
| --- | --- |
| Mouse KPC-1 cells (pdx-1::Cre;KrasG12D;p53R172) | Kind gift from Prof. J. Morton |
| Mouse CKPC-1 and CKPC-2 PDAC cells (from BSNA9.4a male mouse and BSNA15.1a female mouse) | This paper |
| COS-7 cells | ATCC |
| HEK293T | ATCC |

### Experimental models

| Name | Source |
| --- | --- |
| Mouse: Pdx-1::Cre;KrasG12D;p53R172H (KPC) mice | Hingorani et al., 2003 |
| Mouse: Fam49b<sup>fl/fl</sup> | This study |
| CD-1 Nude Mice | Charles River |

### DNA constructs

| Name | Source |
| --- | --- |
| CYRI-B-p17-GFP | Le et al., 2021 |
| pEGFP-N1 | Prof. Laura Machesky lab |
| LPAR1-mCherry | Juin et al., 2019 |
| LPAR1-GFP | Juin et al., 2019 |
| HA-LPAR1 | Kind gift from Dr. Heidi Welch |
| Rab5a-mCherry | Addgene #55126 |

## Supplementary Information

**Supplementary Figure S1:**
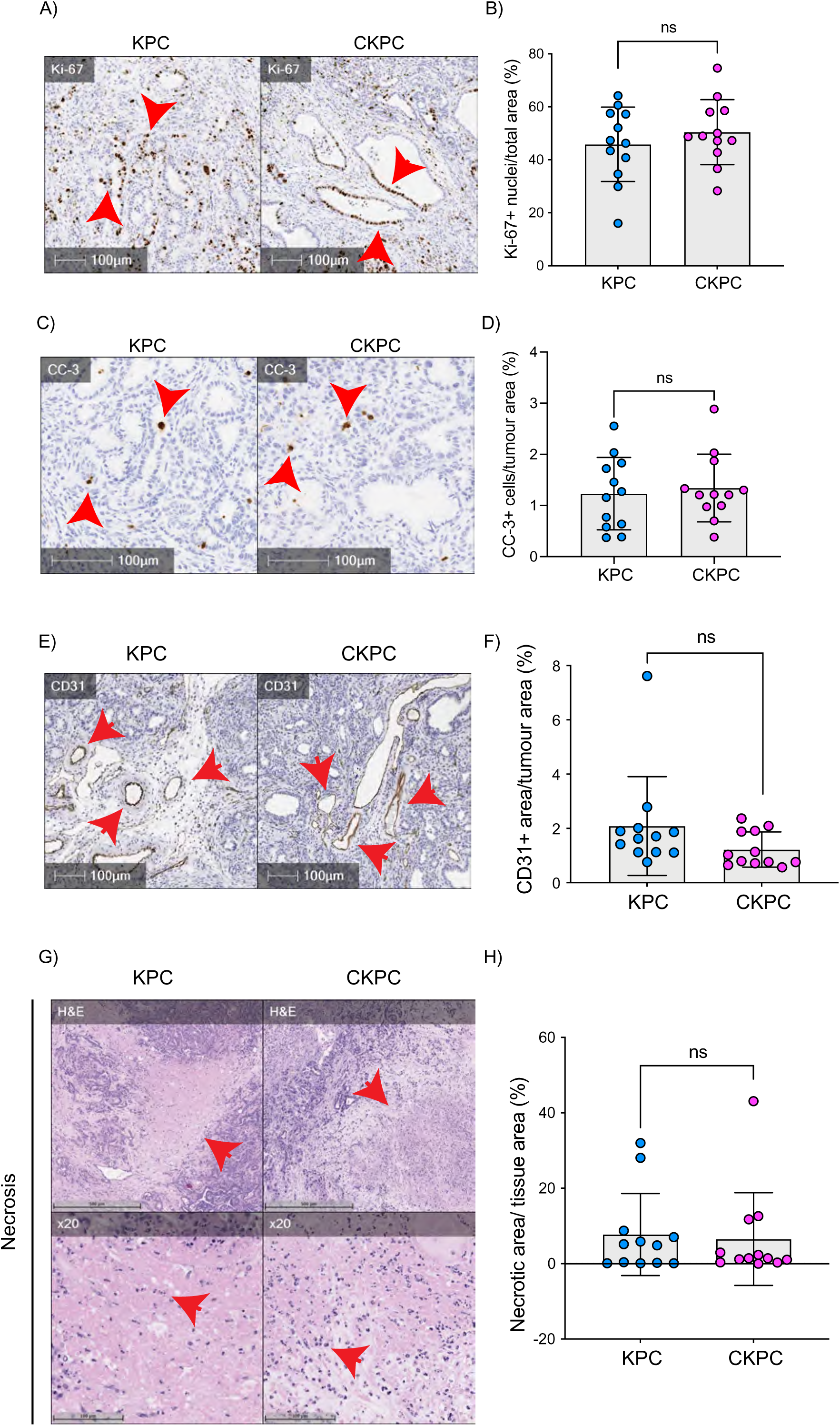
End-point CKPC tumors show comparable proliferation, apoptosis vascularisation and necrosis to KPC tumours. A) Representative images of PDAC sections in KPC and CKPC end-point mice, stained for Ki-67 (proliferation). Scale bars, 100µm. Red arrows indicate the positive cells. B) Quantification of Ki-67 positive nuclei per area from A. Mean ± SD; Unpaired t-test was performed in n=12 KPC and 12 CKPC independent mice. p-value: ns. C) Representative images of PDAC sections in KPC and CKPC end-point mice, stained for cleaved caspase 3 (CC-3, apoptosis). Scale bars, 100µm. Red arrows indicate the positive cells. D) Quantification of CC-3 positive cells per area from c. Mean ± SD; Unpaired t-test was performed in n=12 KPC and 12 CKPC independent mice. p-value: ns. E) Representative images of PDAC sections in KPC and CKPC end-point mice, stained for CD31 (endothelial marker). Scale bars, 100µm. Red arrows indicate the positive area for CD31. F) Quantification of CD31 positive area per tumour area from E. Mean ± SD; Mann-Whitney test was performed in n=12 KPC and 12 CKPC independent mice. p-value: ns = not significant. G) Representative images of PDAC section in KPC and CKPC end-point mice, stained for haematoxylin and eosin (H&E) to identify necrotic areas. Scale bar, 500µm. Red arrows indicate the positive area for necrosis. x20 objective used to show the fragmented nuclei within the necrotic areas (Red arrows). Scale bar, 100µm. H) Quantification of necrotic area per tumour area from G. Mean ± SD; Mann-Whitney test was performed in n=12 KPC and 12 CKPC independent mice. p-value: ns.

**Supplementary Figure S2:**
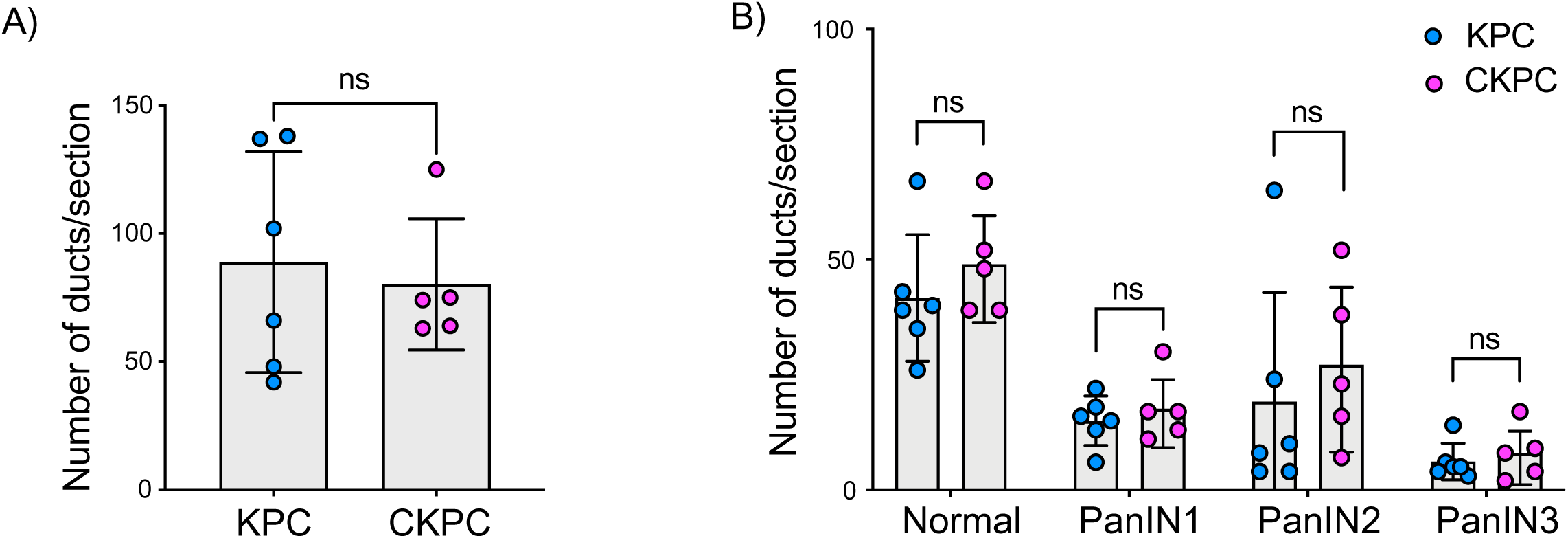
Loss of CYRI-B does not alter the formation of PanIN lesions in 10- week old mice. A) Quantification of the number of ducts present in 10-week-old pancreas in KPC and CKPC mice (n≥5 mic E). Mean ± SD; unpaired t-test was performed. ns=not significant. B) Classification and scoring of pancreatic ducts at in pancreas from 10-week-old KPC and CKPC mice (n≥5 micE). Mean ± SD; unpaired t-test was performed. Ns=not significant.

**Supplementary Figure S3:**
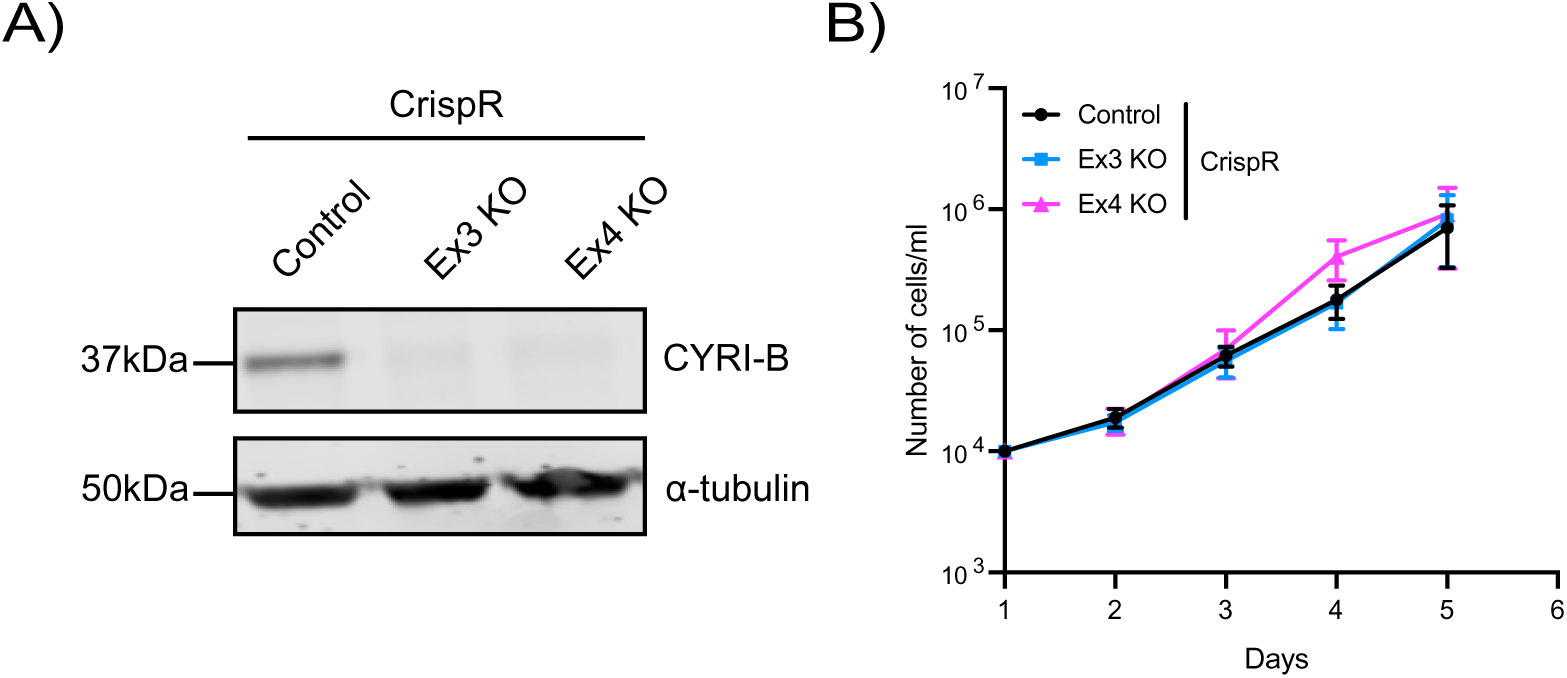
Deletion of CYRI-B in KPC-1 cells does not affect proliferation. A) Representative Western blot from KPC-1 cells for CYRI-B KO. Empty vector was used as control. For CYRI-B KO, Ex3 and Ex4 sequences were used. Alpha-Tubulin was used as loading control. Molecular weights as indicated on the side. B) Proliferation assay of control or CYRI-B KO KPC-1 cells from A). 10^4^ cells were seeded on Day1 and manually counted every day. Error bars represent mean ± SEM from n=3 independent repeats.

**Supplementary Figure S4:**
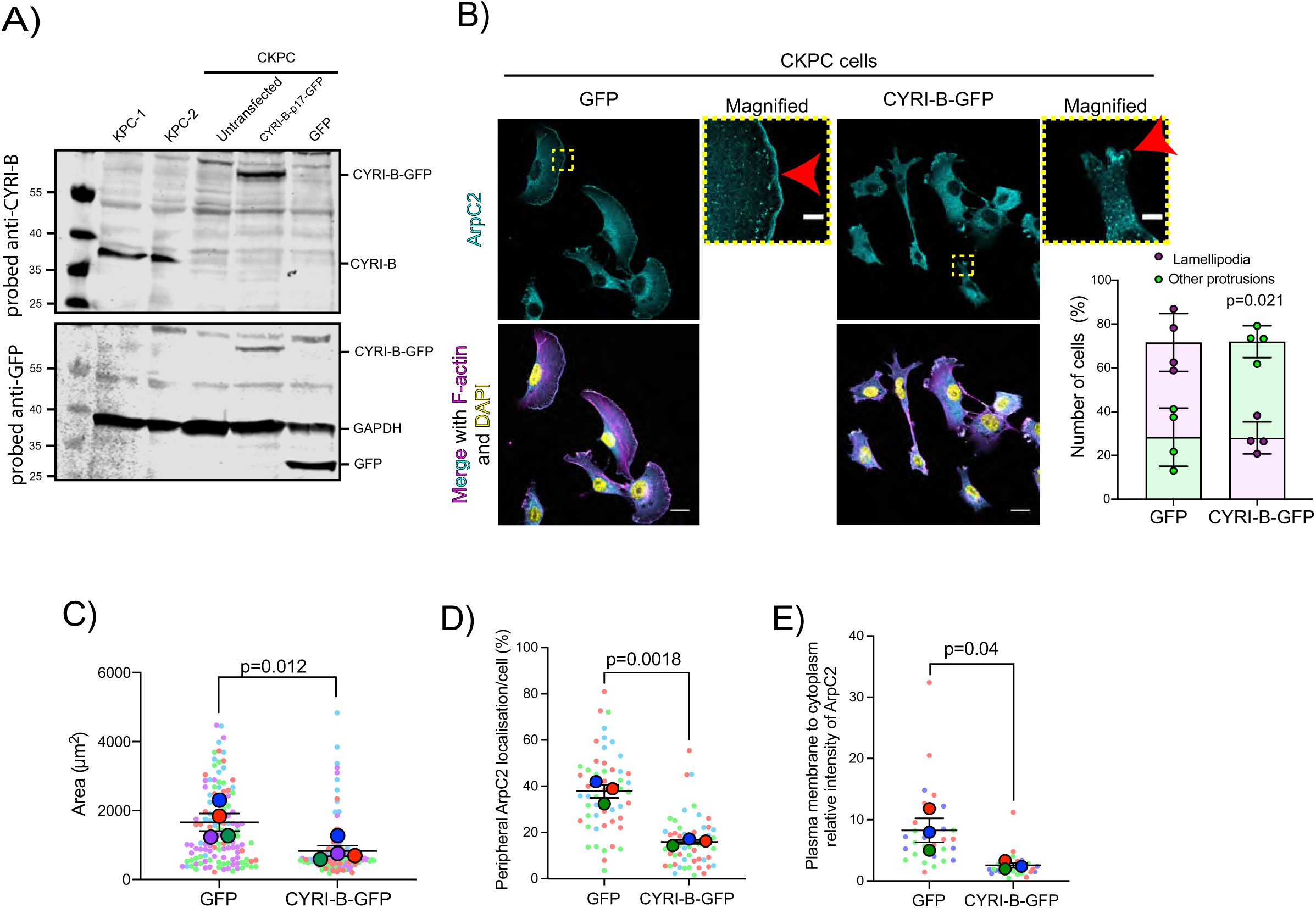
Loss of CYRI-B results in enhanced spreading and Arp2/3 leading edge recruitment in PDAC cells. A) Western blot images of CKPC-1 cells stably expressing CYRI-B-p17-GFP or GFP. KPC-1 and KPC-2 cell lines were used as control. Untransfected CKPC cells were also used as a control. Membranes were probed for anti-GFP (bottom blot) and anti-CYRI-B (top blot). GAPDH was used as loading control. Molecular weights are displayed on the side. B) Representative immunofluorescence images of CKPC CYRI-B KO and rescued cells. Cells were seeded on fibronectin coated coverslips, fixed and stained for F-actin (magenta), ArpC2 (cyan) and DAPI for nuclei (yellow). Scale bars, 20μm. Yellow dotted boxes show the sites for the magnified images. Red arrows show the positive area for ArpC2 staining at the leading edge. Scale bars, 5μm Graph shows manual quantification of the number of cells presenting with lamellipodia (purplE) or other protrusions (green) from B. Mean ± S.D.; Paired t-test was performed in n=4. p-value as indicated. C) Quantification of cell area per cell from B) based on the F-actin staining. Scatter plot here is presented as super plot and every independent biological repeat is coloured differently. Mean ± S.E.M; Paired t-test was performed in n=4. p-value as indicated. D) Manual quantification of the length of the cell periphery showing strong ArpC2 accumulation at, normalised to the total cell periphery. Scatter plot here is presented as a super plot and every independent biological repeat is coloured differently. Mean ± S.E.M; Unpaired t-test was performed in n=3 (from a total of 30 cells). p-value as indicated. E) Manual quantification of the relative intensity of ArpC2 on the plasma membrane to cytoplasmic average intensity. Scatter plot here is presented as a super plot and every independent biological repeat is coloured differently. Mean ± S.E.M; Unpaired t-test was performed in n=3. p-value as indicated.

**Supplementary Figure S5:**
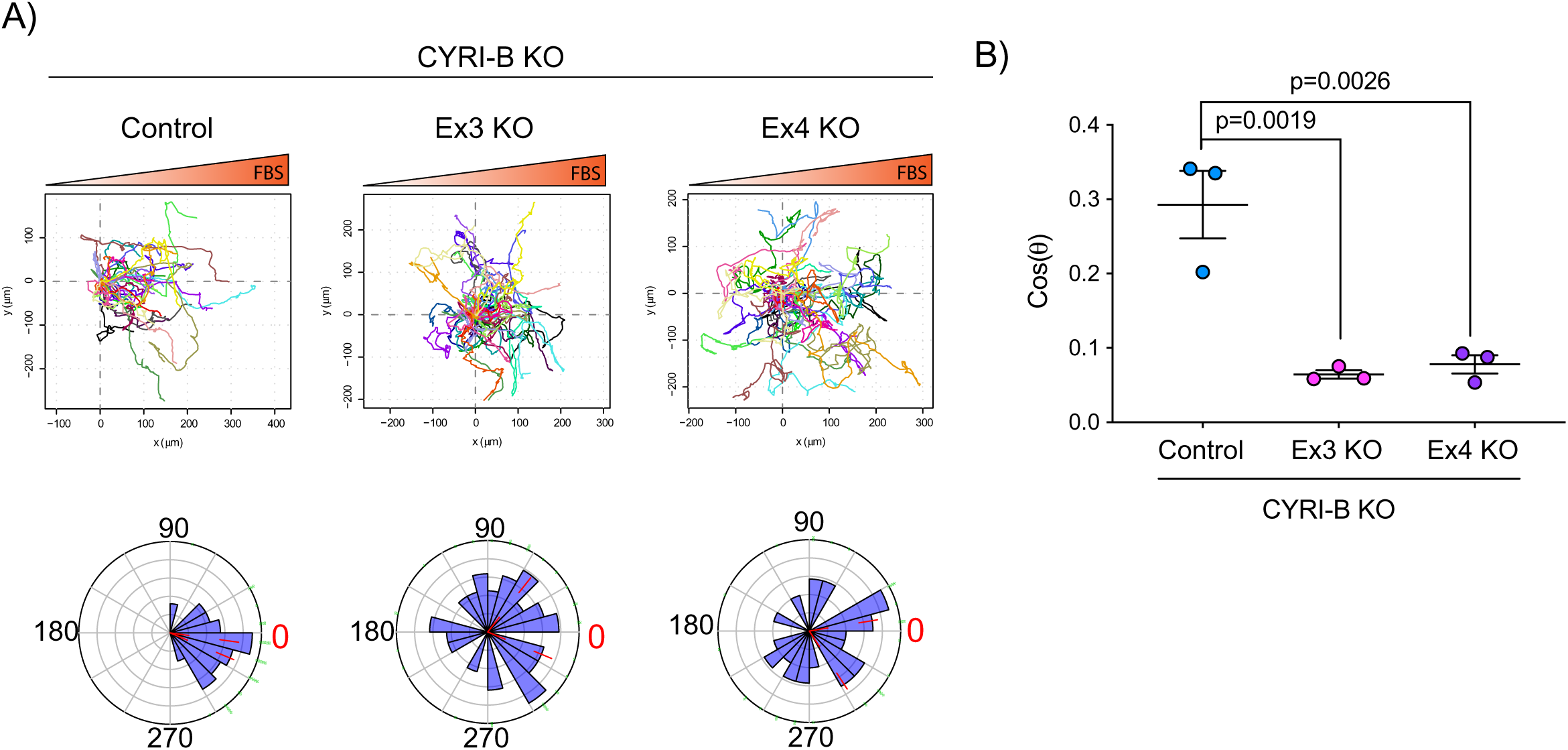
Deletion of CYRI-B abolishes chemotaxis. A) Representative spider-plots from n=3 independent chemotaxis assays of KPC-1 control or CYRI-B KO (EX3 and EX4) cells. Cells were seeded on fibronectin coated coverslips and the “Insall” chamber was assembled. A chemotactic gradient of 10% FBS was established and cells were imaged for 16h (1 frame/15min). Every cell trajectory is displayed with a different colour and the displacement of each cell is reported in the x and y-axis. Orange gradient above shows the FBS gradient. Rose-plot data are displayed for each condition below. Red dashed lines show the 95% confidence interval for the mean direction in the Rose plots. The numbers represent degrees of the angle of migration relative to the chemoattractant, with zero (reD) denoting the direction of the chemoattractant gradient. B) Quantification of the results in A) showing the cos(θ) data (chemotactic index). Mean ± S.E.M from the average cos(θ) data of every repeat; One-Way ANOVA followed by Tukey’s multiple comparisons test was performed. Red dashed lines were indicated show the 95% confidence interval for the mean direction in the Rose plots. p-values as indicated on the graph.

**Supplementary Figure S6:**
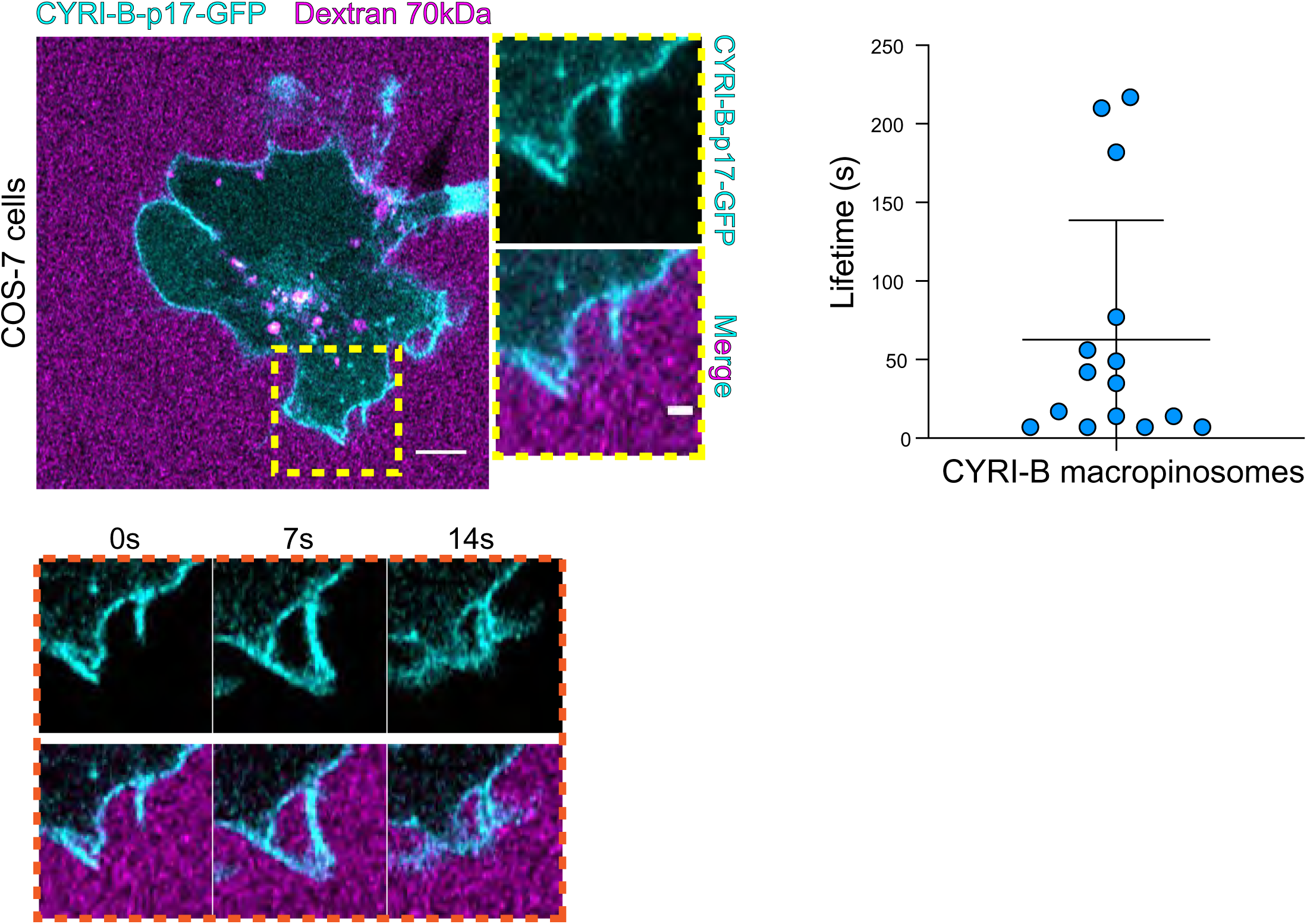
CYRI-B localises at macropinocytic cups in COS-7 cells. Still image from live-cell imaging of COS-7 CYRI-B KO cells transfected with CYRI-B-p17-GFP (cyan). 70kDa Dextran was added to the medium to visualise the macropinocytic events (magenta). Scale bar, 5μm. Yellow box shows the magnified area of interest, showing the macropinocytic cups. Scale bar, 5μm. Scatter plot represents the lifetime of CYRI-B+ macropinosomes once internalised. Mean ± SD. Orange box shows a representative montage of CYRI-B internalisation via macropinocytosis. n=15 events from a total of 6 cells.

**Supplementary Figure S7:**
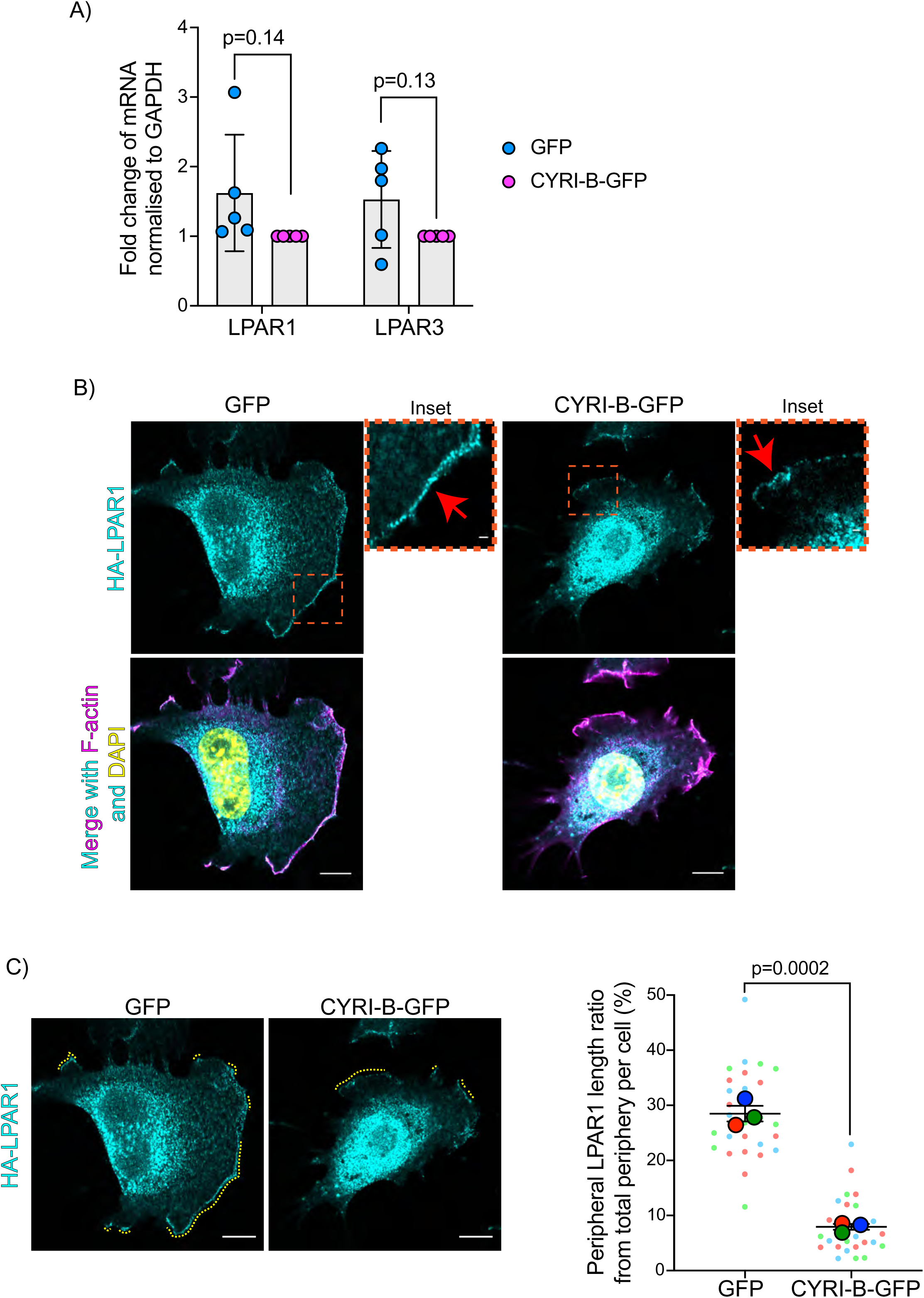
Loss of CYRI-B alters membrane localisation of LPAR1 but not its expression. A) qPCR analysis for endogenous gene expression of LPAR1 and LPAR3 in CKPC-1 stable cells either transfected with GFP or CYRI-B-GFP. The histogram shows the relative mRNA expression from rescued CYRI-B-GFP and normalised from GAPDH expression. Error bars show the mean ± SD. Unpaired t-test was performed in n=5 independent repeats. p-value as indicated. B) Representative immunofluorescence images of CKPC-1 CYRI-B KO and rescued cells. Cells were transfected with LPAR1-HA, then seeded on fibronectin coated coverslips, fixed and stained for F-actin (magenta), anti-HA (cyan) and DAPI for nuclei (yellow). Scale bars, 10μm. Orange dotted boxes show the sites for the magnified images (inset). Red arrows show the positive area for LPAR1 staining at the leading edge. Scale bars, 1μm C) Representative immunofluorescence images from B) as an example of how analysis was performed. Yellow dotted lines show the LPAR1 positive area at the periphery of the cells. Right panel shows the manual quantification of the length of LPAR1 in the periphery of the cells, normalised to the total cell length. Scatter plot here is presented as super plots and every independent biological repeat is coloured differently. Mean ± S.E.M; Unpaired t-test was performed in n=3 (from a total of 28 cells). p-value as indicated. Scale bars, 10μm.

## Supplementary Video and Table legends

**Supplementary Video 1 (for** **Figure 5): CYRI-B is localised at internal vesicles and tubules**

Live-cell video of COS-7 CYRI-B KO cells transfected with CYRI-B-p17-GFP (cyan). Time interval, 5sec. Scale bar, 5μm.

**Supplementary Video 2 (for** **Figure 5): CYRI-B is localised at membrane cups**

Live-cell video of COS-7 CYRI-B KO cells transfected with CYRI-B-p17-GFP (cyan). Time interval, 5sec. Scale bar, 2μm.

**Supplementary Video 3 (for** **Figure 6): CYRI-B is localised at macropinocytic cups and internal macropinosomes in PDAC cells**

Live-cell video of BSNA cells stably transfected with CYRI-B-p17-GFP (cyan). 70kDa dextran (magenta) was added to the medium as a marker of macropinocytosis. Time interval, 5sec. Scale bar, 5μm.

**Supplementary Video 4 (for Supplementary Figure 6): CYRI-B is localised at macropinocytic cups in COS-7 cells**

Live-cell video of COS-7 CYRI-B KO cells transfected with CYRI-B-p17-GFP (cyan). 70kDa dextran (magenta) was added to the medium as a marker of macropinocytosis. Time interval, 7sec. Scale bar, 2μm.

**Supplementary Video 5 (for** **Figure 6): CYRI-B is recruited to macropinocytic cups and precedes Rab5 recruitment**

Live-cell video of COS-7 CYRI-B KO cells transfected with CYRI-B-p17-GFP (cyan) and mRFP-Rab5 (magenta). Time interval, 10sec. Scale bar, 5μm.

**Supplementary Video 6 (for** **Figure 7): LPAR1 internalises via macropinocytosis**

Live-cell video of COS-7 cells transfected with LPAR1-GFP (cyan). 70kDa Dextran was added to the medium to visualise the macropinosomes (magenta). Time interval, 6sec. Scale bar, 2μm.

**Supplementary Video 7 (for** **Figure 7): LPAR1 internalises via CYRI-B positive macropinosomes**

Live-cell video of CKPC-1 cells transfected with CYRI-B-p17-GFP (cyan) and LPAR1-mCherry (magenta). Time interval, 10sec. Scale bar, 5μm.

**Table S1:** Spreadsheet with data on cohort animals. Table outlining data on mouse cohorts used in the experiments.

| Survival curve |  |  |  |  |  |  |  |  |  |  |
| --- | --- | --- | --- | --- | --- | --- | --- | --- | --- | --- |
| Mouse ID | Sex | D.O.B | CRE | CYRI-B | Kras G12D | p53 R172H | Cohort | age, days | D.O.D | Status |
| BSNA1.2a | Male | 19-Jul-18 | pos | wt | het | het | Survival | 126 | 19/11/2018 |  |
| BSNA7.1c | Female | 15-Aug-18 | Pos | wt | het | het | Survival | 424 | 19/10/2019 | censored |
| BSNA7.1e | Female | 19-Aug-18 | Pos | wt | het | het | Survival | 95 | 23/11/2018 | censored |
| BSNA5.1e | Female | 19-Aug-18 | Pos | wt | het | het | Survival | 91 | 19/11/2018 | censored |
| BSNA7.2b | Male | 11-Sep-18 | Pos | wt | het | het | Survival | 72 | 23/11/2018 | censored |
| BSNA7.2d | Female | 11-Sep-18 | Pos | wt | het | het | Survival | 170 | 26/02/2019 |  |
| BSNA7.2g | Female | 11-Sep-18 | Pos | wt | het | het | Survival | 130 | 21/01/2019 |  |
| BSNA13.1e | Female | 13-Sep-18 | Pos | wt | het | het | Survival | 114 | 07/01/2019 | censored |
| BSNA5.2c | Female | 12-Sep-18 | Pos | wt | het | het | Survival | 296 | 08/07/2019 |  |
| BSNA14.1d | Female | 25-Sep-18 | pos | wt | het | het | Survival | 159 | 06/03/2019 | censored |
| BSNA14.1g | Female | 25-Sep-18 | pos | wt | het | het | Survival | 167 | 01/05/2019 |  |
| BSNA12.2b | Male | 11-Oct-18 | Pos | wt | het | het | Survival | 71 | 22/12/2018 | censored |
| BSNA10.3c | Male | 28-Oct-18 | Pos | wt | het | het | Survival | 71 | 09/01/2019 | censored |
| BSNA9.3c | Female | 29-Oct-18 | Pos | wt | het | het | Survival | 95 | 04/02/2019 | censored |
| BSNA15.3a | Male | 13-Nov-18 | Pos | wt | het | het | Survival | 125 | 18/03/2019 | censored |
| BSNA20.1d | Female | 27-Dec-18 | Pos | wt | het | het | Survival | 188 | 05/07/2019 | censored |
| BSNA28.1d | Male | 28-Jan-19 | Pos | wt | het | het | Survival | 102 | 10/05/2019 | censored |
| BSNA31.1b | Male | 27-Feb-19 | Pos | wt | het | het | Survival | 87 | 24/05/2019 | censored |
| BSNA35.1a | Male | 04-Apr-19 | Pos | wt | het | het | Survival | 125 | 09/08/2019 |  |
| BSNA35.1c | Male | 04-Apr-19 | Pos | wt | het | het | Survival | 205 | 29/10/2019 |  |
| BSNA35.1g | Female | 04-Apr-19 | Pos | wt | het | het | Survival | 207 | 01/11/2019 |  |
| BSNA36.1d | Female | 08-Apr-19 | Pos | wt | het | het | Survival | 212 | 10/11/2019 |  |
| BSNA31.3c | Female | 15-Apr-19 | Pos | wt | het | het | Survival | 98 | 22/07/2019 | censored |
| BSNA38.1d | Female | 10-Apr-19 | Pos | wt | het | het | Survival | 152 | 12/09/2019 |  |
| BSNA41.1a | Male | 05-May-19 | Pos | wt | het | het | Survival | 207 | 02/12/2019 |  |
| BSNA41.1d | Female | 05-May-19 | Pos | wt | het | het | Survival | 157 | 12/10/2019 | censored |
| BSNA34.2c | Male | 10-May-19 | pos | wt | het | het | Survival | 180 | 10/11/2019 |  |
| BSNA39.1c | Female | 16-May-19 | pos | wt | het | het | Survival | 86 | 12/08/2019 |  |
| BSNA39.1f | Female | 16-May-19 | pos | wt | het | het | Survival | 120 | 16/09/2019 |  |
| BSNA41.2d | Female | 28-May-19 | pos | wt | het | het | Survival | 117 | 25/09/2019 |  |
| BSNA43.2d | Male | 30-May-19 | Pos | wt | het | het | Survival | 92 | 02/09/2019 |  |
| BSNA32.5b | Male | 01-Jun-19 | Pos | Wt | het | het | Survival | 102 | 13/09/2019 |  |
| BSNA43.3a | Male | 22-Jun-19 | Pos | wt | het | het | Survival | 167 | 09/12/2019 |  |
| BSNA43.3c | Male | 22-Jun-19 | Pos | wt | het | het | Survival | 204 | 16/01/2020 |  |
| BSNA41.3b | Male | 02-Jul-19 | Pos | wt | het | het | Survival | 64 | 06/09/2019 | censored |
| BAJF91.1a | Male | 19-Mar-18 | pos | wt | het | het | Survival | 272 | 21/12/2018 | censored |
| BAJF91.1d | Female | 19-Mar-18 | pos | wt | het | het | Survival | 133 | 26/07/2018 |  |
| BAJF97.1a | Male | 10-Jun-18 | pos | wt | het | het | Survival | 146 | 03/09/2018 | censored |
| BAJF97.1b | Male | 10-Jun-18 | pos | wt | het | het | Survival | 169 | 26/09/2018 | censored |
| BAJF108.1a | Male | 02-May-19 | pos | wt | het | het | Survival | 139 | 20/09/2019 | censored |
| BAJF108.1f | Female | 02-May-19 | pos | wt | het | het | Survival | 170 | 21/10/2019 |  |
| BAJF104.1d | Male | 02-Jun-19 | pos | wt | het | het | Survival | 66 | 08/08/2019 | censored |
| BAJF107.2b | Male | 14-Jun-19 | pos | wt | het | het | Survival | 67 | 21/08/2019 |  |
| BSNA9.1d | Male | 17-Aug-18 | Pos | hom | het | het | Survival | 132 | 04/01/2019 |  |
| BSNA8.1c | Male | 21-Aug-18 | Pos | hom | het | het | Survival | 220 | 01/04/2019 | censored |
| BSNA8.1f | Female | 21-Aug-18 | Pos | hom | het | het | Survival | 118 | 19/12/2018 |  |
| BSNA12.1a | Male | 16-Sep-18 | Pos | hom | het | het | Survival | 103 | 30/12/2018 |  |
| BSNA13.1b | Female | 13-Sep-18 | Pos | hom | het | het | Survival | 99 | 22/12/2018 |  |
| BSNA8.2c | Male | 26-Sep-18 | pos | hom | het | het | Survival | 117 | 23/01/2019 |  |
| BSNA15.1a | Female | 27-Sep-18 | Pos | hom | het | het | Survival | 171 | 18/03/2019 |  |
| BSNA11.1f | Female | 25-Oct-18 | Pos | hom | het | het | Survival | 179 | 24/04/2019 |  |
| BSNA11.1g | Female | 25-Oct-18 | Pos | hom | het | het | Survival | 93 | 28/01/2019 |  |
| BSNA9.4a | Male | 08-Dec-18 | Pos | hom | het | het | Survival | 70 | 18/02/2019 |  |
| BSNA9.4d | Male | 08-Dec-18 | Pos | hom | het | het | Survival | 112 | 30/03/2019 |  |
| BSNA19.1b | Male | 25-Nov-18 | Pos | hom | het | het | Survival | 195 | 10/06/2019 |  |
| BSNA16.1a | Male | 24-Nov-18 | Pos | hom | het | het | Survival | 180 | 24/05/2019 | censored |
| BSNA10.4d | Female | 17-Dec-18 | Pos | hom | het | het | Survival | 97 | 24/03/2019 |  |
| BSNA24.1c | Male | 20-Dec-18 | Pos | hom | het | het | Survival | 55 | 15/02/2019 | censored |
| BSNA24.1d | Male | 20-Dec-18 | Pos | hom | het | het | Survival | 173 | 13/06/2019 |  |
| BSNA11.3a | Male | 22-Dec-18 | Pos | hom | het | het | Survival | 104 | 06/04/2019 |  |
| BSNA33.1c | Male | 27-Feb-19 | Pos | hom | het | het | Survival | 124 | 01/07/2019 | censored |
| BSNA31.2c | Female | 23-Mar-19 | Pos | hom | het | het | Survival | 105 | 08/07/2019 |  |
| BSNA38.1c | Female | 10-Apr-19 | Pos | hom | het | het | Survival | 92 | 12/07/2019 | censored |
| BSNA37.2b | Male | 28-Apr-19 | Pos | hom | het | het | Survival | 156 | 04/10/2019 |  |
| BSNA34.2i | Female | 10-May-19 | pos | hom | het | het | Survival | 159 | 19/10/2019 |  |
| BSNA49.2c | Female | 12-Sep-19 | pos | hom | het | het | survival | 98 | 20/12/2019 |  |
| BSNA56.1g | Female | 04-Nov-19 | pos | hom | het | het | survival | 139 | 23/03/2020 |  |
| BSNA58.1d | Male | 26-Nov-19 | pos | hom | het | het | survival | 100 | 06/03/2020 |  |
| BSNA60.1b | Male | 31-Dec-19 | pos | hom | het | het | survival | 76 | 16/03/2020 |  |
| BSNA64.1h | Female | 02-Jan-20 | pos | hom | het | het | Survival | 91 | 03/04/2020 |  |
| BAJF91.1a | Male | 19-Mar-18 | pos | wt | het | het | Survival | 272 | 21/12/2018 | censored |
| BAJF91.1d | Female | 19-Mar-18 | pos | wt | het | het | Survival | 133 | 26/07/2018 |  |
| BAJF97.1a | Male | 10-Jun-18 | pos | wt | het | het | Survival | 146 | 03/09/2018 | censored |
| BAJF97.1b | Male | 10-Jun-18 | pos | wt | het | het | Survival | 169 | 26/09/2018 | censored |
| BAJF108.1a | Male | 02-May-19 | pos | wt | het | het | Survival | 139 | 20/09/2019 | censored |
| BAJF108.1f | Female | 02-May-19 | pos | wt | het | het | Survival | 170 | 21/10/2019 |  |
| BAJF104.1d | Male | 02-Jun-19 | pos | wt | het | het | Survival | 66 | 08/08/2019 | censored |
| BAJF107.2b | Male | 14-Jun-19 | pos | wt | het | het | Survival | 67 | 21/08/2019 |  |

Early time points
| Mouse ID | Sex | D.O.B | CRE | CYRI-B | Kras G12D | p53 R172H | Cohort | age, days | D.O.D | Status |
| --- | --- | --- | --- | --- | --- | --- | --- | --- | --- | --- |
| BSNA47.1a | Male | 20-Aug-19 | pos | wt | het | het | 15 weeks | 104 | 04/12/2019 |  |
| BSNA54.1d | Female | 07-Nov-19 | pos | wt | het | het | 15 weeks | 107 | 24/02/2020 |  |
| BSNA53.2a | Male | 19-Nov-19 | pos | wt | het | het | 15 weeks | 105 | 04/03/2020 |  |
| BSNA71.1d | Female | 19-Mar-20 | pos | wt | het | het | 15 weeks | 104 | 03/07/2020 |  |
| BSNA80.1g | Female | 30-Apr-20 | pos | wt | het | het | 15 weeks | 795 | 14/08/2020 |  |
| BSNA84.3c | Female | 27-Jul-20 | pos | wt | het | het | 15 weeks | 103 | 10/11/2020 |  |
| BSNA87.2d | Male | 16-Aug-20 | pos | wt | het | het | 15 weeks | 105 | 01/12/2020 |  |
| BSNA77.1c | Female | 29-Apr-20 | pos | wt | het | het | 10 weeks | 69 | 08/07/2020 |  |
| BSNA77.2c | Male | 25-May-20 | pos | wt | het | het | 10 weeks | 770 | 05/08/2020 |  |
| BSNA84.1e | Male | 02-Jun-20 | pos | wt | het | het | 10 weeks | 763 | 10/08/2020 |  |
| BSNA81.1e | Male | 01-Jun-20 | pos | wt | het | het | 10 weeks | 764 | 10/08/2020 |  |
| BSNA88.1b | Female | 16-Aug-20 | pos | wt | het | het | 10 weeks | 70 | 26/10/2020 |  |
| BSNA88.1h | Female | 16-Aug-20 | pos | wt | het | het | 10 weeks | 70 | 26/10/2020 |  |
| BSNA66.1j | Female | 30-Jan-20 | pos | wt | het | het | 6 weeks | 42 | 12/03/2020 |  |
| BSNA87.1b | Male | 23-Jul-20 | pos | wt | het | het | 6 weeks | 712 | 03/09/2020 |  |
| BSNA87.1e | Male | 23-Jul-20 | pos | wt | het | het | 6 weeks | 712 | 03/09/2020 |  |
| BSNA87.1g | Male | 23-Jul-20 | pos | wt | het | het | 6 weeks | 712 | 03/09/2020 |  |
| BSNA87.1k | Female | 23-Jul-20 | pos | wt | het | het | 6 weeks | 712 | 03/09/2020 |  |
| BSNA67.1a | Male | 30-Jan-20 | pos | hom | het | het | 6 weeks | 42 | 12/03/2020 |  |
| BSNA66.1g | Female | 30-Jan-20 | pos | hom | het | het | 6 weeks | 42 | 12/03/2020 |  |
| BSNA66.1i | Female | 30-Jan-20 | pos | hom | het | het | 6 weeks | 42 | 12/03/2020 |  |
| BSNA84.3d | Female | 27-Jul-20 | pos | hom | het | het | 6 weeks | 708 | 07/09/2020 |  |
| BSNA84.3e | Female | 27-Jul-20 | pos | hom | het | het | 6 weeks | 708 | 07/09/2020 |  |
| BSNA63.3a | Female | 03-Mar-20 | pos | hom | het | het | 10 weeks | 69 | 12/05/2020 |  |
| BSNA68.2a | Male | 03-Mar-20 | pos | hom | het | het | 10 weeks | 69 | 12/05/2020 |  |
| BSNA69.3c | Female | 17-Mar-20 | pos | hom | het | het | 10 weeks | 69 | 26/05/2020 |  |
| BSNA80.1a | Male | 30-Apr-20 | pos | hom | het | het | 10 weeks | 68 | 08/07/2020 |  |
| BSNA89.1e | Female | 04-Sep-20 | pos | hom | het | het | 10 weeks | 69 | 13/11/2020 |  |
| BSNA64.1e | Male | 02-Jan-20 | pos | hom | het | het | 15 weeks | 105 | 17/04/2020 |  |
| BSNA64.1f | Female | 02-Jan-20 | pos | hom | het | het | 15 weeks | 105 | 17/04/2020 |  |
| BSNA60.2h | Female | 23-Jan-20 | pos | hom | het | het | 15 weeks | 103 | 06/05/2020 |  |
| BSNA70.2c | Male | 23-Feb-20 | pos | hom | het | het | 15 weeks | 105 | 08/06/2020 |  |
| BSNA69.3a | Male | 17-Mar-20 | pos | hom | het | het | 15 weeks | 104 | 01/07/2020 |  |
| BSNA80.1f | Female | 30-Apr-20 | pos | hom | het | het | 15 weeks | 795 | 14/08/2020 |  |
| BSNA90.2e | Male | 12-Oct-20 | pos | hom | het | het | 15 weeks | 105 | 26/01/2021 |  |

RNAscope
| Mouse ID | Sex | D.O.B | CRE | CYRI-B | Kras G12D | p53 R172H | Cohort | age, days | D.O.D | Status |
| --- | --- | --- | --- | --- | --- | --- | --- | --- | --- | --- |
| BAJD34.1h | Male | 07-Dec-16 | pos | wt | het | het | 6 weeks | 42 | 19/01/2017 |  |
| BAJD92.2d | Female | 15-Oct-17 | pos | wt | het | het | 6 weeks | 45 | 30/11/2017 |  |
| BAJD97.1a | Male | 19-Oct-17 | pos | wt | het | het | 6 weeks | 38 | 27/11/2017 |  |
| BAJD91.1d | Female | 21-Sep-17 | pos | wt | het | het | 10 weeks | 69 | 30/11/2017 |  |
| BAJD91.1f | Female | 21-Sep-17 | pos | wt | het | het | 10 weeks | 69 | 30/11/2017 |  |
| BAJD92.1b | Male | 21-Sep-17 | pos | wt | het | het | 10 weeks | 66 | 27/11/2017 |  |
| BAJD91.2b | Female | 01-Nov-17 | pos | wt | het | het | 10 weeks | 69 | 10/01/2018 |  |
| BAJD82.2a | Male | 26-Jul-17 | pos | wt | het | het | 15 weeks | 102 | 08/11/2017 |  |
| BAJD90.1d | Male | 26-Jul-17 | pos | wt | het | het | 15 weeks | 102 | 08/11/2017 |  |
| BAJD93.1b | Male | 23-Sep-17 | pos | wt | het | het | 15 weeks | 102 | 05/01/2018 |  |
| BAJD93.1d | Female | 23-Sep-17 | pos | wt | het | het | 15 weeks | 102 | 05/01/2018 |  |
| BAJD91.2d | Female | 01-Nov-17 | pos | wt | het | het | 15 weeks | 103 | 14/02/2018 |  |
| BAJD81.2b | Male | 08-Jun-17 | pos | wt | het | het | End point | 75 | 23/08/2017 |  |
| BAJD25.2a | Male | 13-Sep-16 | pos | wt | het | het | End point | 177 | 10/03/2017 |  |
| BAJD35.1j | Female | 28-Oct-16 | pos | wt | het | het | End point | 180 | 28/04/2017 |  |
| BAJD14.1a | Male | 12-Sep-16 | pos | wt | het | het | End point | 205 | 07/04/2017 |  |

**Bedding** Datasand Eco-Pure Aspen Chips 4 Premium **Sizzle-Pet** LBS Biotechnology **Dome Home** LBS Biotechnology **Plastic tunnel tube** LBS Biotechnology **Enrichment** Datasand Aspen Bricks 500x100x100mm **Diet** Social Diet Services CRME irradiated **Cages** Techniplast GM500
D.O.B. = Date of birth, D.O.D. = Date of death, pos = positive for allele, het = heterozygous, wt = wild type

## Notes

### Competing Interest Statement

The authors have declared no competing interest.

